# Disordered region of nuclear membrane protein Bqt4 recruits phosphatidic acid to the nuclear envelope to maintain its structural integrity

**DOI:** 10.1101/2023.12.21.572941

**Authors:** Yasuhiro Hirano, Tsukino Sato, Ayane Miura, Yoshino Kubota, Tomoko Shindo, Koichi Fukase, Tatsuo Fukagawa, Kazuya Kabayama, Tokuko Haraguchi, Yasushi Hiraoka

## Abstract

The nuclear envelope (NE) is a permeable barrier that maintains nuclear–cytoplasmic compartmentalization and ensures nuclear function; however, it ruptures in various situations such as mechanical stress and mitosis. Although the protein components for sealing a ruptured NE have been identified, the mechanism by which lipid components are involved in this process remains to be elucidated. Here, we found that an inner nuclear membrane (INM) protein Bqt4 directly interacts with phosphatidic acid (PA) and serves as a platform for NE maintenance in the fission yeast *Schizosaccharomyces pombe*. The intrinsically disordered region (IDR) of Bqt4, proximal to the transmembrane domain, binds to PA and forms a solid aggregate *in vitro*. Excessive accumulation of Bqt4 IDR in INM results in membrane overproliferation and lipid droplet formation in the nucleus, leading to centromere dissociation from the NE and chromosome missegregation. Our findings suggest that Bqt4 IDR controls nuclear membrane homeostasis by recruiting PA to the INM, thereby maintaining the structural integrity of the NE.

## Introduction

In eukaryotic cells, the nuclear envelope (NE) encapsulates chromosomes in the nucleus. The NE functions as a permeable barrier to maintain nuclear–cytoplasmic compartmentalization and ensures various nuclear functions. The NE comprises inner and outer nuclear membranes (INM and ONM, respectively), which comprise lipid membranes and transmembrane proteins. The INM interacts with chromosomes to regulate their organization and genetic activity. The ONM is continuous with the endoplasmic reticulum (ER). The integrity of the NE, which retains the nuclear barrier, is crucial for cell viability (Webster and Lusk, 2016), and yet, the NE is dynamic and undergoes repeated breakdown and reformation during mitosis (open mitosis) in higher eukaryotic cells (Güttinger et al., 2009; Lajoie and Ullman, 2017; Ungricht and Kutay, 2017). The NE also ruptures occasionally because of nuclear damage, such as mechanical stress, during the interphase (Denais et al., 2016). Thus, the ruptured NE should be promptly resealed to ensure nuclear functions. Recent studies have identified the components essential for NE sealing; however, the underlying mechanisms have not been fully understood.

To reseal incidental NE ruptures, eukaryotic cells have a mechanism that surveils and maintains the NE integrity. To maintain the NE integrity, the endosomal sorting complex required for transport-III (ESCRT-III) is transiently recruited to the rupture sites of the NE by charged multivesicular body protein 7 (CHMP7/Chm7/Cmp7 in metazoans, budding yeast *Saccharomyces cerevisiae*, and fission yeast *Schizosaccharomyces pombe* and *japonicus*) (Gu et al., 2017; Olmos et al., 2015; Pieper et al., 2020; Vietri et al., 2015; Von Appen et al., 2020). Lem2, a member of the conserved LEM (Lap2β-Emerin-Man1) domain INM protein family, functions as a nuclear adaptor for CHMP7 (Gu et al., 2017; Von Appen et al., 2020; Webster et al., 2016). ESCRT-III seals membrane rupture sites during mitosis and interphase (Denais et al., 2016; Raab et al., 2016). Recent studies have shown that this sealing mechanism is highly conserved and plays a global role in diverse eukaryotes (Dey et al., 2020; Gu et al., 2017).

In addition to the proteinaceous sealing factors, the NE rupture sealing mechanism involves *de novo* lipid synthesis (Bahmanyar and Schlieker, 2020; Kinugasa et al., 2019; Lee et al., 2020; Penfield et al., 2020). Glycerophospholipids, major components of cellular lipid membranes, are essential for nuclear membrane expansion (Siniossoglou, 2013; Foo et al., 2023; Takemoto et al., 2016). The synthesis of glycerophospholipids in the NE is regulated by the phosphatidate phosphatase lipin (Ned1 in *S. pombe*; Pah1 in *S. cerevisiae*), CNEP1 (an NE-localizing activator of lipin, known as the Nem1-Spo7 complex in yeast), and phosphatidate cytidylyltransferase Dgk1 in metazoans (Han et al., 2006; Romanauska and Köhler, 2018). In *C. elegans* embryos, the CNEP1-lipin pathway coordinates the production of ER membranes, which narrow the holes in the NE ruptures (Bahmanyar, 2015; Bahmanyar et al., 2014; Penfield et al., 2020). Additionally, ESCRT-III complexes limit excess ER membrane supplements at these holes (Penfield et al., 2020). These findings suggest that glycerophospholipid synthesis is regulated during NE rupture sealing. Furthermore, the very-long-chain fatty acid synthase Elo2 rescues the NE rupture phenotype in a Lem2 and Bqt4 double deletion mutant in *S. pombe* (Kinugasa et al., 2019), and long-chain sphingoid bases suppress NE morphology defects (Hwang et al., 2019). Although the lipid components required for NE maintenance have been identified, the mechanisms by which these lipids accumulate and function in the INM remains unclear.

Several hundred INM-specific proteins, such as LEM proteins and lamin B receptor, have been identified in mammalian cells (Korfali et al., 2012). INM proteins are involved in NE maintenance and potentially regulate nuclear membrane properties. In *S. pombe*, the INM proteins, Lem2 and Bqt4, play central roles in maintaining NE integrity. Loss of either Lem2 or Bqt4 does not affect cell viability, but loss of both confers a synthetic lethal phenotype accompanied by nuclear membrane rupture (Kinugasa et al., 2019; Tange et al., 2016), suggesting that these proteins share vital functions in NE rupture sealing. Lem2 is a multifunctional protein involved in many cellular processes such as gene expression and NE shaping (Banday et al., 2016; Barrales et al., 2016; Hirano et al., 2020; Kume et al., 2019; Martín Caballero et al., 2022; Tange et al., 2016). Among the wide range of Lem2 functions, the key function of Lem2 for NE maintenance is the recruitment of ESCRT-III to the NE (Gu et al., 2017; Lee et al., 2020; Pieper et al., 2020; Thaller et al., 2021; Von Appen et al., 2020). In contrast to Lem2, the role of Bqt4 in NE maintenance has not been identified. Bqt4 was originally identified as a protein that anchors telomeres to the NE (Chikashige et al., 2009), but the telomere-anchoring function of Bqt4 is unnecessary for NE maintenance (Tange et al., 2016). Our recent studies have shown that excess Bqt4 accumulation in INM deforms the NE (Le et al., 2023) and Bqt4 interacts with Cwh43, an enzyme involved in triacylglycerol metabolism (Hirano et al., 2023a), suggesting that Bqt4 may regulate the properties of INM.

In this study, we investigated the role of Bqt4 in NE maintenance using several techniques: viability assays, lipid binding assays, and imaging techniques including fluorescence live cell imaging, fluorescence recovery after photobleaching (FRAP), fluorescence correlation spectroscopy (FCS), and correlative light-electron microscopy (CLEM).

## Results

### Vital Bqt4 domain required for cell growth

To reveal a Bqt4 function in NE maintenance, we investigated functional domains of Bqt4 using the *lem2*-shut-off *bqt4*Δ strain, which confers synthetic lethality upon shut-off by the addition of thiamine (Hirano et al., 2023b; Kinugasa et al., 2019; Tange et al., 2016). Bqt4 fragments were introduced into this strain and cell viability was tested under shut-off conditions. Based on the annotations in PomBase, we divided Bqt4 into three domains, as illustrated in the schematic diagram (Fig. 1A): 1-140 amino acid (aa) residues containing APSES domain (named for Asm1p, Phd1p, Sok2p, Efg1p, and StuAp); 141-262 aa residues containing a coiled-coil region; and 259-383 aa residues referred to as the “homologous to apolipoprotein” region. GFP-tagged Bqt4 fragments deleted for each of these domains (Δ1-140, Δ141-262, and Δ259-383) were expressed in the *lem2*-shut-off *bqt4*Δ cells and growth of the cells were observed. Upon shut-off of *lem2*, the cells expressing Δ1-140 and Δ259-383 fragments of Bqt4 did not grow, whereas the cells expressing Δ141-262 did (Fig. 1B). These results indicate that 1-140 and 259-383 aa domains are essential for vital functions of Bqt4. All these fragments as well as wild-type Bqt4 (FL) were localized to the NE (Fig. 1C), suggesting that the lethality was not caused by the loss of NE localization.

**Figure 1.**
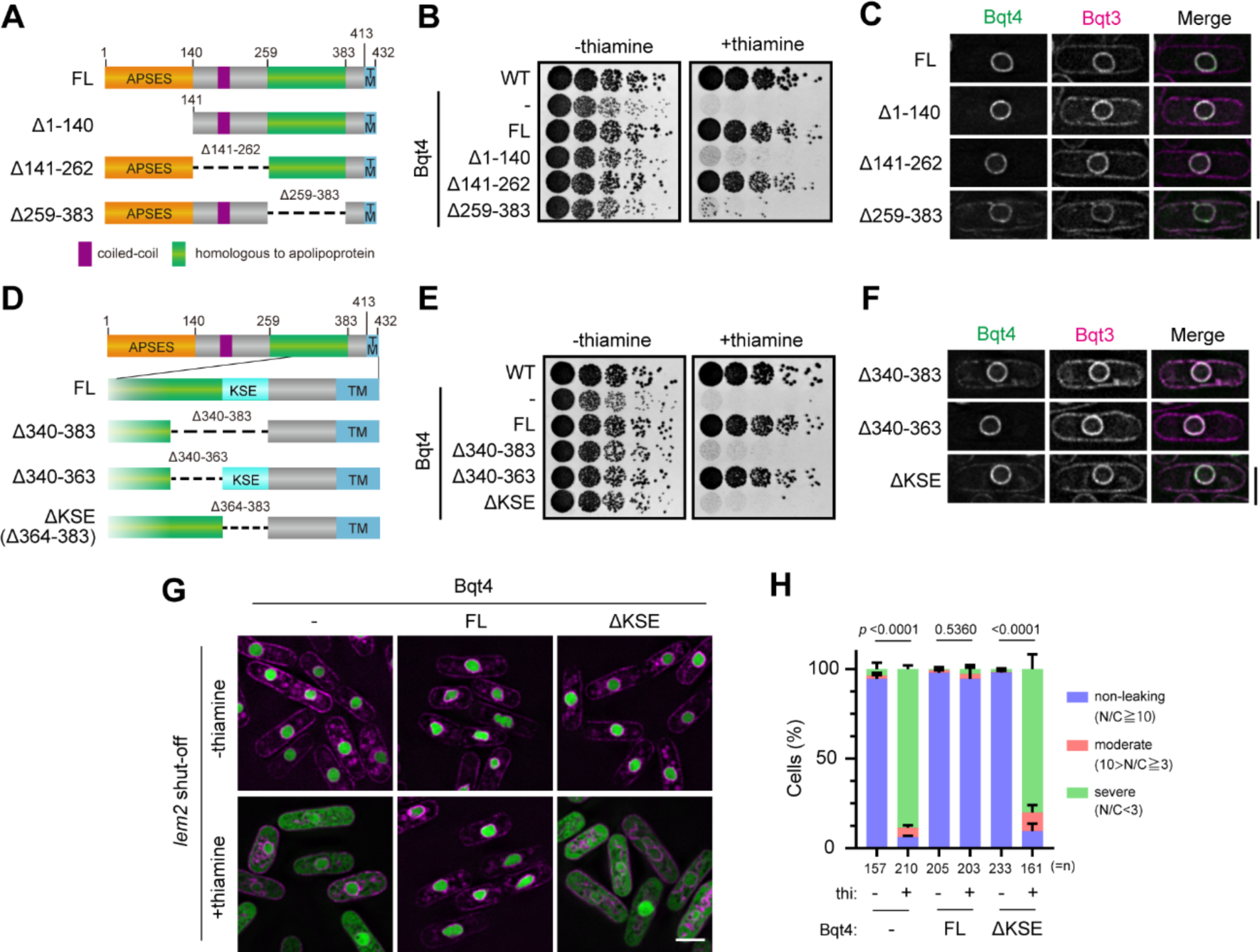
Vital domains for Bqt4 function. **A.** Schematic diagrams of the Bqt4-deletion mutants. Orange, purple, green, and sky-blue squares denote APSES, coiled-coil, homologous to apolipoprotein, and transmembrane (TM) domains, respectively. **B.** Spot assay of Bqt4-deletion mutants (shown in A) as indicated on the left (full-length (FL), Δ1-140, Δ141-262 and Δ259-383; minus (-) indicates GFP only). The GFP-tagged Bqt4 mutants were expressed in the *lem2* shut-off *bqt4*Δ cells, which are synthetic lethal upon thiamine addition. WT represents the wild-type *S. pombe* strain. Five-fold serially diluted cells were spotted on EMMG plates in the absence (-thiamine) or presence (+thiamine) of thiamine, and growth of these cells was observed after 3 days. **C.** Localization of the Bqt4 fragments shown in A. GFP-Bqt4 and mCherry-Bqt3 were co-expressed and observed. The scale bar represents 5 μm. **D.** Schematic diagrams of Bqt4-deletion mutants at amino acid residues 340-383. The 364-383 aa region was named KSE. **E.** Spot assay of the Bqt4-deletion mutants shown in D. GFP-Bqt4 fragments indicated on the left were expressed in the *lem2* shut-off *bqt4*Δ cells, and the growth of these cells was observed as described in B. **F.** Localization of the Bqt4 fragments shown in D. GFP-Bqt4 fragments and mCherry-Bqt3 were co-expressed and observed. The scale bar represents 5 μm. **G.** Nuclear protein leakage in cells expressing ΔKSE fragment. The *lem2* shut-off *bqt4*Δ cells expressing FL or ΔKSE fragment were cultured in the presence or absence of thiamine and GFP–GST–NLS (nuclear protein marker, green) and Ish1–mCherry (NE marker, magenta) signals were observed using fluorescence microscopy. The scale bar represents 5 μm. **H.** Quantification of the nuclear protein leakage phenotype. Fold enrichment of GFP– GST–NLS signal in the nucleus (nucleus/cytoplasm; N/C) was quantified in three levels according to its leakage: non-leaking (N/C≧10), moderate (10>N/C≧3), and severe (N/C<3). Percentages of the cells were obtained from three independent experiments and the mean with s.d. were plotted as a 100% stacked bar chart. *n* indicates total cell number counted. The statistical significance was assessed using the Chi-square test.

The 259–383 aa region was annotated as “homologous to apolipoprotein.” Apolipoprotein is a co-factor protein that binds to lipoproteins, such as low-density lipoprotein and high-density lipoprotein comprising cholesterol and triacylglyceride particles covered with phospholipids, and mediates their transport and metabolism (Mahley et al., 1984). Considering its properties, this region potentially binds to lipids and is consequently involved in NE maintenance; however, this potential interaction has not been experimentally demonstrated. Thus, in this study, we focused on the 259–383 aa region.

### KSE domain of Bqt4 is vital for NE maintenance

To narrow down the vital domain in the 259-383 aa region, we predicted the secondary structure of Bqt4 using the PSIPRED 4.0 prediction server (http://bioinf.cs.ucl.ac.uk/psipred/; Supplementary Fig. 1A). From this prediction, we examined the 340-383 aa region which consists of α-helix (340-363 aa) and low complex regions (364-383 aa) (Fig. 1D and red and blue squared regions in Supplementary Fig. 1A). The deletion mutants Δ340-383 and Δ364-383, but not Δ340-363, showed growth defect in the *lem2*-shut-off *bqt4*Δ strain (Fig. 1E), indicating that the 364-383 aa region possesses vital functions of Bqt4. The 364-383 aa region is biased in positive charge, containing seven positive (K or R), six polar (S), and four negative (E or D) amino acids out of the 20 amino acids (Supplementary Fig. 1A), hereafter referred to as the KSE domain. The ΔKSE mutant as well as wild-type Bqt4 localized at the NE (Fig. 1F) and retained interaction with Bqt3 (Supplementary Fig. 1B) and the telomere anchoring (Supplementary Fig. 1C). Thus, this function is probably distinct from the known Bqt4 functions.

We next examined whether the KSE domain is required for NE maintenance. The ΔKSE mutant was expressed in *lem2-*shut-off *bqt4*Δ cells expressing GFP–GST– NLS, a nuclear protein marker, and nuclear protein leakage from the nucleus was assessed by calculating the fluorescence intensity ratio of nucleus to cytoplasm. Upon thiamine addition, GFP–GST–NLS severely leaked from the nucleus of ΔKSE mutant-expressing cells, but not from the nucleus of FL-expressing cells (Fig. 1G and H), indicating NE rupture in the ΔKSE mutant-expressing cells. These results suggest that the KSE domain is vital for NE maintenance.

### KSE-containing intrinsically disordered region (IDR) domain is responsible for dominant-negative effect of Bqt4 overexpression

We have previously reported that abnormal Bqt4 accumulation causes severe NE deformation-associated growth defects (Le et al., 2023). As the KSE domain is involved in NE maintenance, it may be responsible for this phenotype. To test this hypothesis, GFP-tagged ΔKSE (Δ364-383 aa) or KSE (364-383 aa) protein was overexpressed in *S. pombe* cells under the *nmt1* promotor. Overexpression of the ΔKSE fragment did not cause the growth defects (Supplementary Fig. S2A-C) and NE deformation (Supplementary Fig. S2D and E), suggesting that the KSE region causes dominant-negative effects of Bqt4 overexpression; however, the KSE region alone did not cause these defects (KSE, Supplementary Fig. S2C), indicating that the KSE region is essential, but not sufficient, for the dominant-negative effects of Bqt4 overexpression. A larger fragment containing the KSE region is necessary for the dominant-negative effects.

Thus, we further searched for domains responsible for the growth defects associated with severe NE deformation upon Bqt4 overexpression. For this purpose, we examined the three-dimensional protein structure around the KSE domain predicted by Alphafold2 (https://alphafold.ebi.ac.uk/entry/O60158) and selected an unstructured region (364-413 aa; Fig. 2A) containing the KSE domain as a candidate. We termed this region as an IDR. The IDR fragment of Bqt4 tagged with GFP (GFP-IDR) was localized to the NE despite the absence of TM and nucleoplasm under suppressive expression conditions (Fig. 2B, +thiamine). In contrast, GFP-KSE was localized to nucleoplasm but not to the NE (Fig. 2C). When GFP-IDR was overexpressed in *S. pombe* cells under the control of the *nmt1* promoter, its overexpression resulted in severe growth defects in the spot assay (Fig. 2D). These results indicate that the overexpression of IDR, but not KSE, confers dominant-negative effects on cell growth and suggest that the IDR domain is substantial for NE maintenance, causing loss-of-function and dominant-negative effects. We subsequently focused on the IDR domain. The essentiality of the IDR domain for NE maintenance could not be assessed because deleting the 384–413 aa region caused protein degradation (Supplementary Fig. S3).

**Figure 2.**
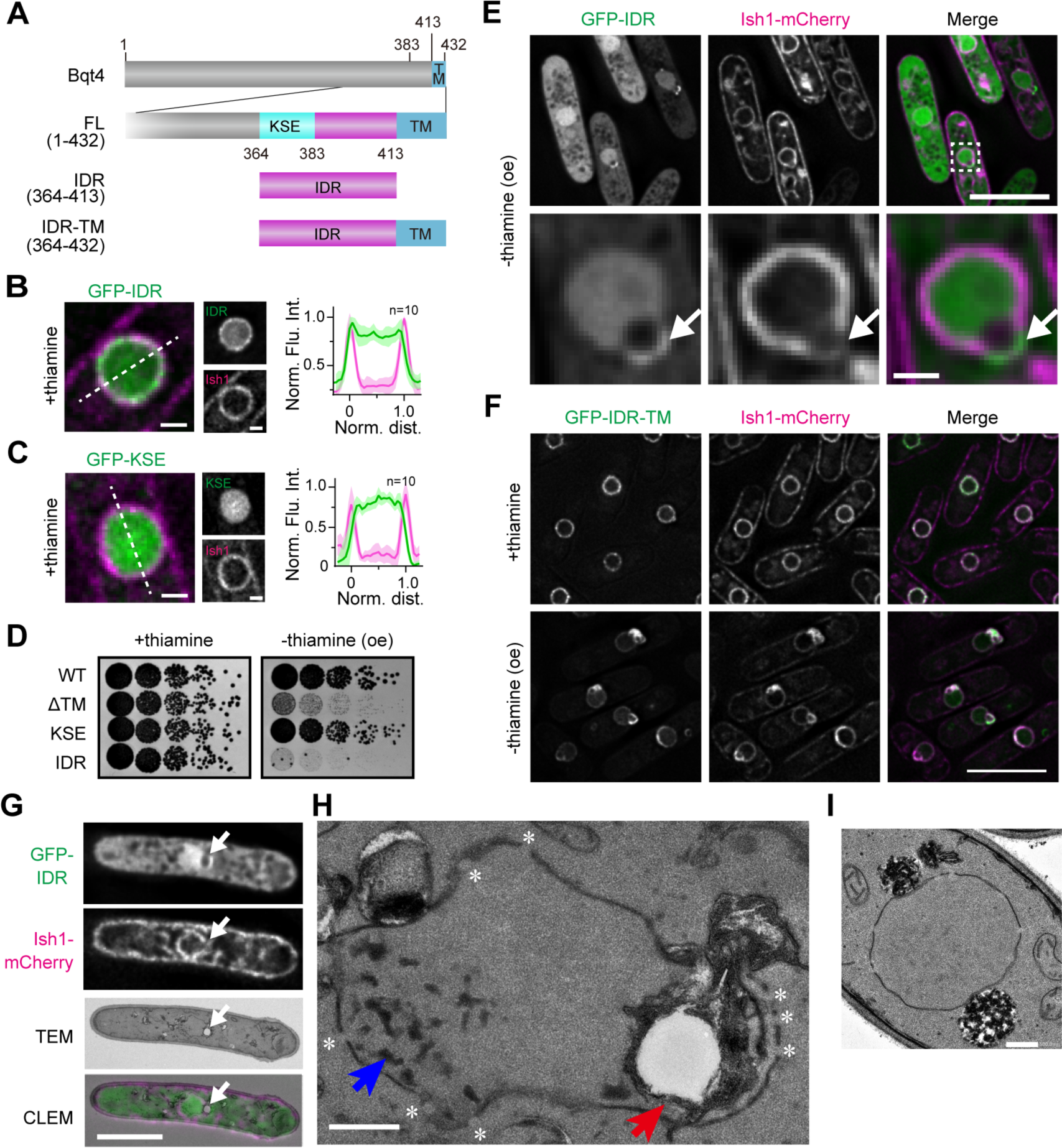
The IDR domain is responsible for maintaining NE structure. **A.** Schematic diagrams of IDR fragments of Bqt4. **B-C**. Localization of IDR (B) and KSE (C) fragments. GFP-IDR or KSE fragments (green) were expressed with Ish1-mCherry (magenta). The cells were cultured in the presence of thiamine and then observed using fluorescence microscopy (left panels). The scale bars represent 1 μm. Quantifications of their localization are shown on the right. Fluorescence intensities of Bqt4 fragments and Ish1 on the white-dashed line were quantified. The normalized mean values calculated from 10 nuclei were plotted with standard deviation along the normalized nuclear size — 0 and 1 represent the NE at both sides of the nucleus. **D**. Spot assay of the cells overexpressing the IDR fragment. GFP-Bqt4 fragments indicated on the left were expressed under the *nmt1* promoter. The growth of the cells was observed as described in Fig. 1B. oe represents overexpression. **E.** Localization of overexpressed IDR. GFP-IDR (green) was overexpressed in cells with Ish1-mCherry (magenta) and the localization was observed using fluorescence microscopy. The dashed square region is enlarged at the bottom. The scale bars in the top and bottom panels represent 5 and 1 μm, respectively. Arrows indicate the herniated NE. **F.** Localization of IDR-TM. GFP-IDR-TM (green) was expressed in cells with Ish1-mCherry (magenta). The cells were cultured in EMMG with (+thiamine) or without (- thiamine (oe)) thiamine and the localization of IDR-TM was observed using fluorescence microscopy. The scale bar represents 5 μm. **G-I**. Representative CLEM images of cells overexpressing IDR. (G) Cells overexpressing IDR were observed using fluorescence microscopy (top 2 frames), followed by transmission electron microscopy (TEM; the third frame). The corresponding CLEM image is shown at the bottom. White arrows in the fluorescence image indicate a herniated NE. Scale bar, 5 μm. (H) A magnified TEM image showing ultrastructures around the nucleus in the same cell shown in (G). Red arrows point to a vacuole-like structure surrounded by a multi-layer membrane, and blue arrows highlight electron-dense patches. Asterisks indicate NPC. Scale bar, 500 nm. (I) TEM image of wild-type cells for comparison. Scale bar, 500 nm.

### Overexpression of IDR induces aberrant NE morphology

Under overexpression conditions, GFP-IDR was localized in the nucleoplasm, accompanied by a herniated structure adjacent to the nucleus (Fig. 2E, arrows). Ish1-mCherry, a marker of NE (Asakawa et al., 2022), was faintly localized in the herniated region, suggesting that the herniated region is somewhat atypical for NE. The Ish1 signal, which was localized in the NE under normal conditions, was distributed on cytoplasmic membranes besides the NE under overexpression conditions (Fig. 2E), suggesting that overexpression of GFP-IDR may cause abnormal membrane proliferation. To confirm the direct effect of IDR overexpression on the NE, we examined the GFP-IDR fragment with a transmembrane domain (IDR-TM; 364-432 aa) and found that overexpression of GFP-IDR-TM caused more severe defects than GFP-IDR in the NE structure, such as herniation (Fig. 2F). These results suggest that the NE localization of overexpressed IDR causes severe NE defects.

To further investigate the effects of IDR on NE structure, we observed cells overexpressing IDR with correlative light-electron microscopy (CLEM) (Fig. 2G and H). The atypical structures observed under live fluorescence imaging (Fig. 2E) were preserved without any obvious alterations during fixation for CLEM (Supplementary Fig. 4A). We found two major abnormalities: a vacuole-like structure covered with a multilayered membrane within the nucleus, and electron-dense patches in the nucleoplasm (Fig. 2H, red and blue arrows, respectively), both of which were not observed in wild-type cells (Fig. 2I). The vacuole-like structure corresponded to the hole observed in the fluorescence image (Fig. 2G, white arrows). In the herniated region, where the IDR signal was concentrated, multilayered membranes extended to the nuclear membrane, and these membranes surrounded the vacuoles. Such abnormal vacuoles in the nucleus were located in close proximity to the NE, which is rich in nuclear pore complexes (NPCs) (Fig. 2H, asterisks and Supplementary Fig. 4 for serial sectioning images). Notably, the nuclear protein did not leak out of the nucleus (Supplementary Fig. 5), indicating that the NE retained its function as a barrier between the nucleus and the cytoplasm despite its abnormal morphology. The electron-dense patches were distributed over a wide area within the nucleoplasm but were scarcely distributed in the nucleolus (see serial sectioning images in Supplementary Fig. 4B). Patches were observed in 45 of 50 nuclei overexpressing IDR. These results suggest that the IDR domain of Bqt4 plays a role in maintaining the NE structure via its membrane proliferation function.

### Overexpression of IDR induces aberrant chromosome organization

The NE is an important architecture for chromosome organization in the nucleus. Therefore, we investigated whether aberrant NE proliferation induced by Bqt4 affects chromosome organization, in which centromeres form a cluster on the NE in *S. pombe* cells. Overexpression of the GFP-IDR and GFP-IDR-TM fragments, but not GFP and GFP-KSE, caused dissociation of the centromeres from the NE and their declustering (Fig. 3A and B). This dissociation did not affect transcriptional silencing in the centromeric region (Supplementary Fig. 6A). The Bqt4-binding protein, Lem2, which binds to the central core of centromeres and is involved in centromere anchoring to the NE, played a minor role in dissociation (Supplementary Fig. 6B and C). Time-lapse observation of the cells overexpressing IDR-TM revealed that aberrant nuclear membrane expansion occurred just after mitosis; subsequently, the clustered centromeres started to dissociate from the NE and then declustered (Fig. 3C and Supplementary Movie 1). Consequently, the chromosome was missegregated during the next mitosis (Fig. 3C). Alternatively, the entire chromosome moved to one side or the cells showed a *cut* phenotype (untimely torn cell phenotype); these cells displayed nuclear membrane extrusion along the spindle microtubule (Supplementary Movie 2). These phenotypes were similar to those of *ned1*-deficient cells; Ned1, a homolog of human lipin, is a phosphatase that generates diacylglycerol (DAG) from PA, and the *ned1-1 ts* mutant induces abnormal membrane proliferation (Tange et al., 2002). Taken together, these observations suggest that NE overproliferation induced by IDR accumulation causes genome instability.

**Figure 3.**
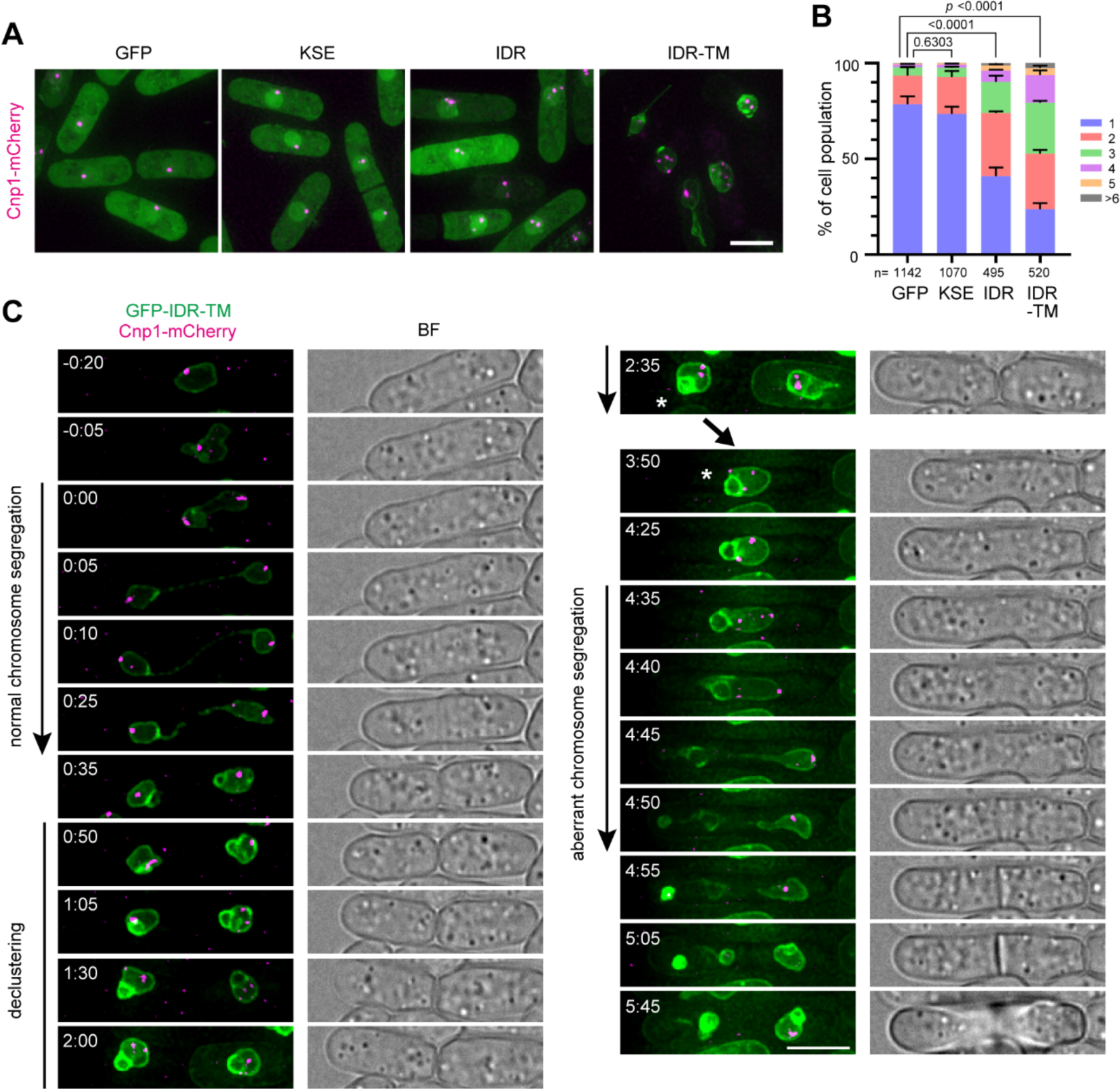
Overexpression of the IDR region disturbs centromere clustering. **A**. Subcellular localization of centromeres in cells overexpressing the IDR fragments. The CENP-A homolog Cnp1 was expressed as an mCherry-tagged protein (magenta) in cells overexpressing Bqt4 fragments, and its localization was observed using fluorescence microscopy. Scale bar represents 10 μm. **B**. Quantification of centromere number. The number of centromeres in each nucleus was counted and plotted as a 100% stacked bar chart. The statistical significance was assessed using the Chi-square test. **C.** Time-lapse imaging. Cells harboring Cnp1-mCherry and *nmt1*-driven GFP-IDR-TM were cultured in EMMG supplemented with 1 μM YAM2 for overnight. After washing out YAM2, the cells were observed using microscopy every 5 min. Elapsed times (h:min) are relative to the onset of first mitosis. The cell labeled by an asterisk (Time 2:35) is shown for the second mitosis (from 3:50 to 5:45). BF denotes a bright-field image. The scale bar represents 5 μm.

### Bqt4 directly binds to PA via the IDR region

The IDR region may be a potential lipid-binding domain because its accumulation in the NE disrupts the membrane structure. To test this hypothesis, we performed a lipid binding assay using a membrane lipid strip spotted on 15 different lipid types (Fig. 4A). Purified His-GFP-tagged IDR fragments were incubated with the membrane, and bound IDR fragments were detected. The IDR fragment was bound to PA and cardiolipin (Fig. 4A, IDR). In this study, we focused on PA in the following experiments because the abnormal membrane proliferation observed with Bqt4 overexpression was similar to that observed in the *ned1*-deficient phenotype. To confirm the binding of PA to the IDR, we performed a liposome-mediated binding assay. Liposomes with different percentages of PA were prepared and incubated with purified GFP-IDR protein. After isolation of the liposomes via density gradient centrifugation, the proteins bound to the liposomes were electrophoresed and detected using silver staining. GFP-IDR co-sedimented with PA-containing liposomes but not with control liposomes (Fig. 4B).

**Figure 4.**
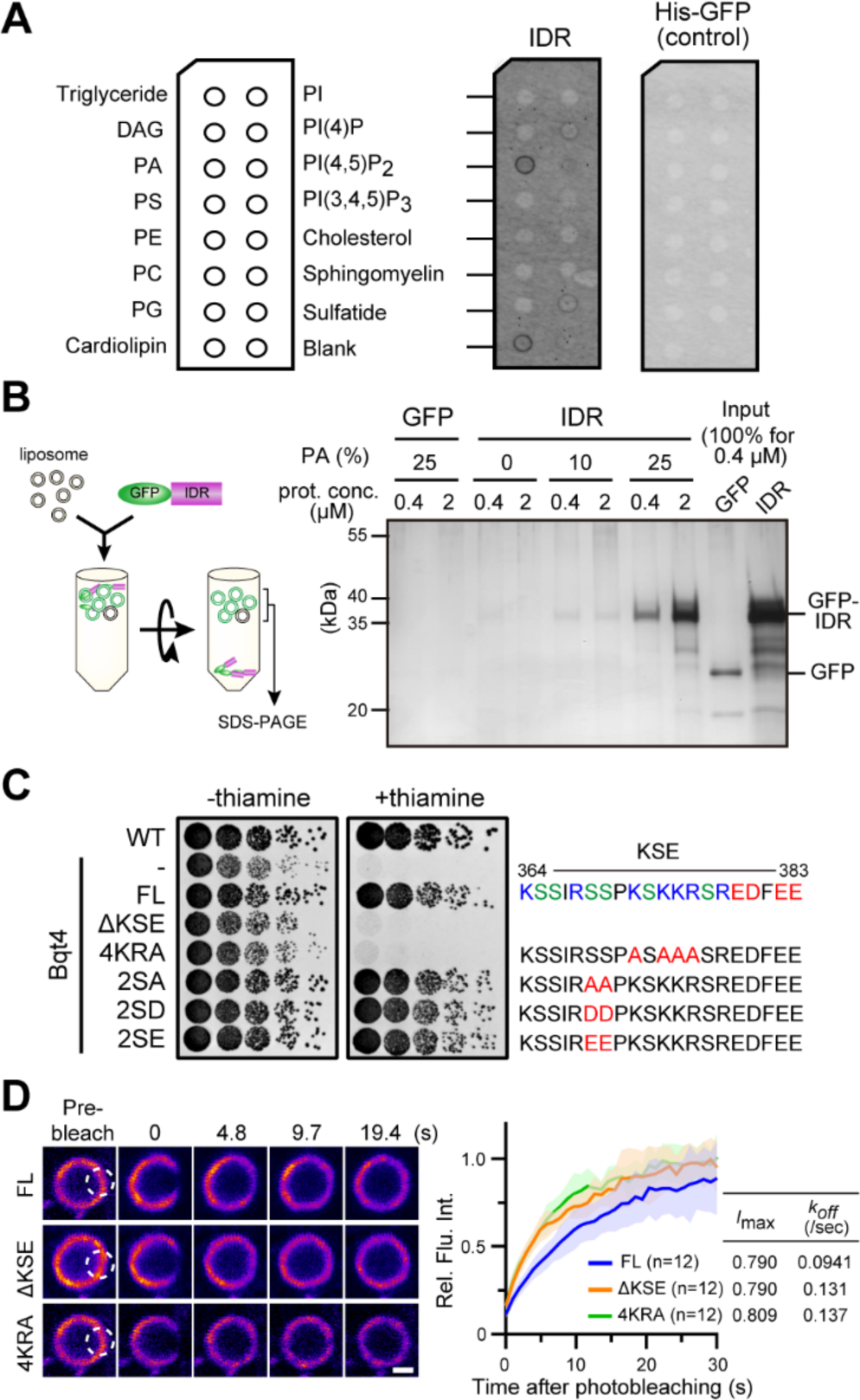
Bqt4 directly binds to PA via IDR. **A**. Lipid binding assay. The schematic illustration on the left shows lipids spotted on the membrane lipid strip. His-GFP-IDR or His-GFP was incubated with the membrane and the bound protein was detected using fluorescence. **B**. Liposome-binding assay. The left diagram shows the procedure of the assay. Liposomes generated from a defined percentage of phospholipids were mixed with GFP or GFP-IDR and then separated via density gradient centrifugation. The protein-liposome complex visualized through light scattering was collected and subjected to SDS-PAGE. Protein components were detected using silver staining. The bound proteins are shown in the right panel. **C**. Spot assay of the point mutants in the KSE region. The cells expressing Bqt4 point mutants shown on the left were spotted and the growth was observed as described in Fig. 1B. The mutation sites in the KSE region are shown on the right. **D**. FRAP analysis of GFP-tagged Bqt4 fragments (FL, ΔKSE and 4KRA). The dashed white open circle region was bleached, and the recovery of fluorescence intensity after bleaching in the circle was measured. Solid lines in the FRAP curve indicate the mean value. The areas on both sides of the solid lines indicate standard deviation. The curves were fitted as a single exponential association, and the maximum recovery rate (*I*_max_) and dissociation coefficient (*k*_off_) are shown on the right. The scale bar represents 1 μm.

Therefore, we conclude that the IDR region directly binds to PA *in vitro*. Because PA and cardiolipin are negatively charged lipids (Fig. 4A), and the IDR region, especially the KSE region, is biased toward a positive charge (Fig. 4C), we assumed that the interaction between PA and the IDR region is mediated via electrostatic interactions. To examine the contribution of these positive amino acids to PA binding, we introduced mutations into the positive amino acids in the KSE region, where four lysine and arginine residues were substituted with alanine (4KRA), and expressed GFP-Bqt4-4KRA in the *lem2* shut-off stain. In addition, we substituted two polar amino acid residues proximal to the positively charged portion with non-polar or negative amino acids (2SA, 2SD, and 2SE) as controls. As predicted, the 4KRA mutant did not restore lethality upon shut-off, whereas the other three mutants did (Fig. 4C), supporting our hypothesis that the IDR and PA electrostatically interact with each other. Next, we examined whether the lipid binding of Bqt4 affects its dynamics in INM using fluorescence recovery after photobleaching (FRAP) (Fig. 4D). The FRAP analysis revealed that GFP-ΔKSE and GFP-4KRA moved faster than full-length Bqt4 (GFP-FL) in the INM, suggesting that Bqt4 forms a sub-stable complex with the INM by interacting with PA. To further confirm the electrostatic interaction between the IDR and PA, we generated mutants by scrambling and reversing the amino acid sequences in the KSE region to disturb its electrostatic potential (Supplementary Fig.7A). These mutants exhibited weak interactions with PA (Supplementary Fig.7B), resulting in a loss of Bqt4 function (Supplementary Fig.7C), and increased mobility in the NE (Supplementary Fig.7D) similar to the 4KRA mutant. This observation supports the notion of electrostatic interactions between the IDR and PA. In addition to the highly positive charge, the position of the sequence was crucial for the IDR-PA interaction, as the reversed fragment lost its functionality. Thus, unlike other inner nuclear membrane proteins, Bqt4 appears to possess the unique property of interacting with PA.

Taken together, these results suggest that Bqt4 binds to PA via electrostatic interactions in the IDR region and that this binding is crucial for Bqt4 function.

### IDR region forms solid-fibrous aggregate in the presence of PA *in vitro*

Based on the predicted structural aspects of the IDR region, we assumed that this region underwent phase separation in the presence of PA. To test this hypothesis, we conducted an *in vitro* condensation assay. As protein concentration is an important factor in analyzing phase separation, we determined the concentration of GFP-Bqt4 in the NE using purified GFP as a standard; GFP-Bqt4 was integrated into the *lys1*^+^ locus and expressed under its own promoter. The protein concentration of Bqt4 in the NE under authentic expression level was estimated to be approximately 1 μM (Supplementary Fig. 8A). In order to understand the phenomenon caused by the abnormal accumulation of the IDR region, we used 2 μM, twice the normal amount, in further experiments. Purified GFP-IDR protein was mixed with an increased concentration of PA, and the mixture was observed using fluorescence microscopy. As shown in Fig. 5A and B, GFP-IDR formed fibrous aggregates in a PA-dependent manner. Consistent with the lipid binding assay results, aggregates were not formed with the neutral phospholipid phosphatidylcholine (PC).

**Figure 5.**
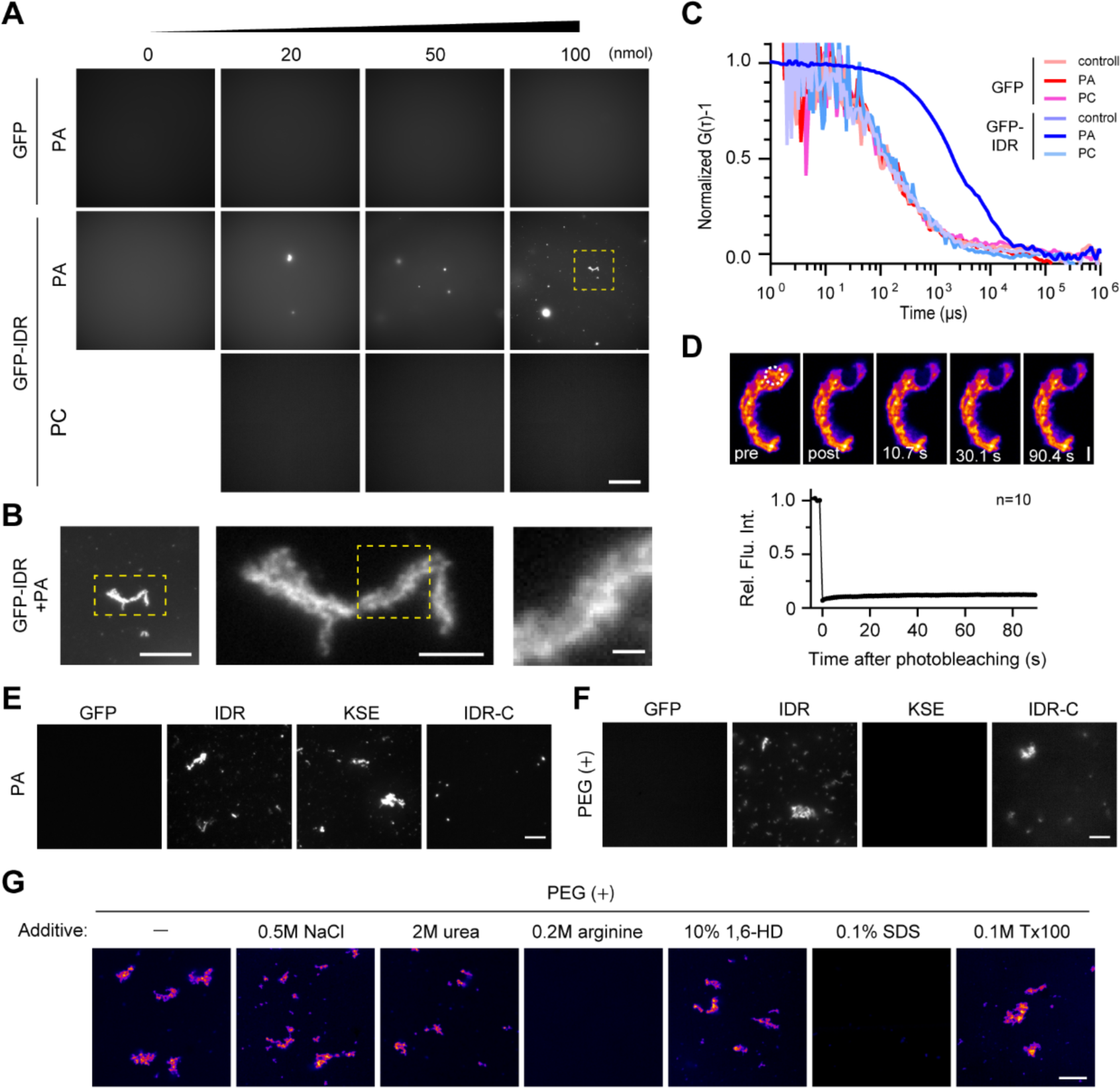
IDR forms solid aggregate with PA. **A-B**. Aggregate formation of GFP-IDR with PA. Purified GFP and GFP-IDR were mixed with increased amounts (0, 20, 50, and 100 nmol) of PA or PC. The aggregates formed were observed using fluorescence microscopy (A). The yellow-squared region is enlarged in (B). The scale bar in (A) represents 50 μm. Scale bars in (B) represent 20, 5, and 1 μm from left to right, respectively. **C.** Molecular clustering state of GFP-IDR aggregates. GFP (left) and GFP-IDR (right) were mixed with 100 nmol PA or PC, and then the mixture was analyzed using FCS. Count rate and normalized auto-correlation function calculated from the measurement (G(τ)) are shown in (C) and (D), respectively. Inlets in (C) are enlarged images of the part of the count rate. **D.** Molecular exchange rate in GFP-IDR aggregate. GFP-IDR was mixed with 100 nmol PA, and the molecular exchange rate of the formed aggregate was measured using FRAP. The white-dashed circle denotes the bleached region. The scale bar represents 2 μm. The fluorescence recovery curve shows mean ± s.e.m. from ten measurements. **E.** Responsible region of aggregate formation in the IDR with PA. GFP, GFP-IDR (IDR), GFP-KSE (KSE), and GFP-IDR-C (IDR-C) were mixed with 100 nmol PA, and the formed aggregate was observed using fluorescence microscopy. The scale bar represents 10 μm. **F.** Responsible region of aggregate formation in the IDR under crowding conditions. GFP, GFP-IDR (IDR), GFP-KSE (KSE), and GFP-IDR-C (IDR-C) were incubated in a crowding buffer containing 5% PEG as a crowder, and the formed aggregate was observed using fluorescence microscopy. The scale bar represents 10 μm. **G.** Effect of additives on the aggregate formation. GFP-IDR was incubated in a crowding buffer supplied with additives indicated on the top, and the formed aggregate was observed using fluorescence microscopy. The scale bar represents 10 μm.

To further assess the aggregation state of GFP-IDR, we performed fluorescence correlation spectroscopy (FCS) analysis. In the presence of PA, fluorescence intensity fluctuations of approximately 100 kHz were accompanied by strong spike signals for the GFP-IDR (Supplementary Fig. 9A), most likely originating from large fibrous aggregates observed in Fig. 5A and B passing through the confocal volume. Conversely, in the absence of PA, fluctuations were less than 10 kHz (Supplementary Fig. 9A, compare inlets for GFP-IDR+PA and control), a range similar to that observed when using GFP instead of GFP-IDR, or PC instead of PA (Supplementary Fig. 9A). The auto-correlation function (G(τ)), calculated from these fluctuations, showed that GFP-IDR exhibits slower diffusion with PA compared to other combinations (Fig. 5C). These results suggest that GFP-IDR formed microscopically invisible clusters with PA, in addition to large visible aggregates. As the large aggregates were not spherical, they are unlikely to have been formed by liquid-liquid phase separation. We also performed a FRAP assay on GFP-IDR within the aggregates and found that its molecular exchange rate was low (Fig. 5D), suggesting that the aggregates are of a solid phase, not a liquid phase. Collectively, the IDR tended to form a cluster in the presence of PA; ultimately, this cluster transitioned into a large, solid, fibrous aggregate *in vitro*.

To explore the key properties of aggregate formation, we divided the IDR into two fragments, GFP-tagged KSE (364-383 aa) and IDR-C (384-413 aa), and evaluated their aggregation-forming activity in the presence of PA. GFP-KSE exhibited marked aggregation-forming activity, whereas GFP-IDR-C showed much lower activity than FL and KSE (Fig. 5E). As shown by the FRAP assay (Fig. 4D), the positive charges in the KSE region restricted Bqt4 dynamics in the NE, likely due to interactions with PA. Nevertheless, IDR-C should have a positive role in the membrane-association activity of IDR because the activity of IDR was stronger than that of KSE (compare Fig. 2B and C).

To further test the functional differences between the KSE and IDR-C regions, we incubated GFP-KSE and GFP-IDR-C in a molecular crowding solution containing polyethyleneglycol (PEG) and assessed their ability to form aggregates. For this experiment, we selected a protein concentration of 8 μM because of the following reasons: 1) the ease of observation via a fluorescence microscope (Supplementary Fig. 9B) and 2) this concentration is in a range below the overexpression under the *nmt1* promoter (Supplementary Fig. 8B). Contrary to the results of PA-mediated aggregate formation (Fig. 5E), GFP-IDR-C exhibited aggregate formation similar to that of the IDR region in the presence of PEG, whereas GFP-KSE did not (Fig. 5F). This result suggests that the IDR-C region can facilitate aggregate formation once the KSE region forms a cluster with PA and increases the local concentration of Bqt4 in the NE.

Finally, we tested the effect of additives on GFP-IDR aggregate formation under crowding conditions to elucidate the dominant interaction type for aggregate formation. Sodium chloride (NaCl), urea, arginine, 1,6-hexanediol (1,6-HD), sodium dodecyl sulfate (SDS), and Triton X-100 (Tx-100) were used as salt, protein denaturant, protein aggregation suppressor, aliphatic alcohol, ionic detergent, and non-ionic detergent, respectively. Arginine and SDS drastically inhibited the aggregate formation among the additives tested (Fig. 5G). These two additives also inhibited the aggregate formation of the IDR-C fragment (Supplementary Fig. 10A). As arginine suppresses the hydrophobic interactions of proteins by binding to the aromatic group, thus increasing the net charge of the protein (Miyatake et al., 2016), the hydrophobic interaction between IDRs is likely a dominant force in aggregation in *in vitro* condition. This result was consistent with the finding that PA-induced aggregate formation because PA could strengthen the hydrophobic interaction between the IDRs by neutralizing the positive charge in the IDR, especially in the KSE region. These observations were consistent with the molecular properties of the IDR region; the KSE and IDR-C regions were hydrophilic and hydrophobic, respectively (Supplementary Fig. 10B and C). However, because the aggregates were formed in the presence of NaCl, 1,6-HD, and Tx-100, multivalent interactions might be involved in the aggregate formation of the IDRs. The IDR-derived aggregates were soluble in low concentrations (0.1%) of SDS, suggesting that the aggregates possess a similar property to fused in sarcoma (FUS) and hnRNPA2 (Kato et al., 2012) and might be able to revert to a dispersion state in the cell, although they are quite stable.

Taken together, these results demonstrate that the IDR region has a property to form aggregates with PA, which is accelerated by the two distinct properties of KSE and IDR-C regions.

### Bqt4 is involved in regulating the balance of lipid synthesis by accumulating PA in the INM

Next, we investigated the biological significance of the interaction between IDR and PA. Given our assumption that IDR overexpression changes the intracellular localization of PA, we generated *S. pombe* cells expressing an mCherry-tagged PA sensor that comprises a homodimer of the PA-binding helix of *S. cerevisiae* Opi1 bearing a G120W point mutation (Foo et al., 2023) to visualize PA localization. This sensor was primarily localized in the nucleus and peripheral ER (Fig. 6A). In the cells overexpressing FL, IDR-TM and IDR of Bqt4, intracellular localization of the PA sensor was dramatically changed (Fig. 6A and B for quantification). In cells overexpressing FL, the PA sensor accumulated in round structures beneath the NE that were likely lipid droplets (LDs) covered with PA-containing phospholipids (Fig. 6A, arrowheads in FL). These droplets were observed near the blebs of Bqt4 signals in the NE. Similar PA accumulation in the nucleus was observed in IDR-TM–overexpressing cells (IDR-TM in Fig. 6A and B for quantification). These results suggested that Bqt4 collects PA-enriched membranes to the NE via its IDR domain. In IDR-overexpressing cells, GFP-IDR was localized in both the nucleoplasm and cytoplasm (Fig. 6A, IDR). Consequently, the PA sensor was redistributed from the nucleus to the cytoplasm (Fig. 6A; IDR). In contrast, KSE overexpression did not alter the localization of the PA sensor, although this fragment was localized in the cytoplasm, similar to IDR (Fig. 6A, KSE). Therefore, we conclude that IDR, but not KSE, controls intracellular PA localization depending on its localization and protein levels in *S. pombe* cells. As the nuclear membrane over-proliferates following Bqt4 overexpression (Fig. 2), PA recruitment possibly accelerates phospholipid synthesis in the NE.

**Figure 6.**
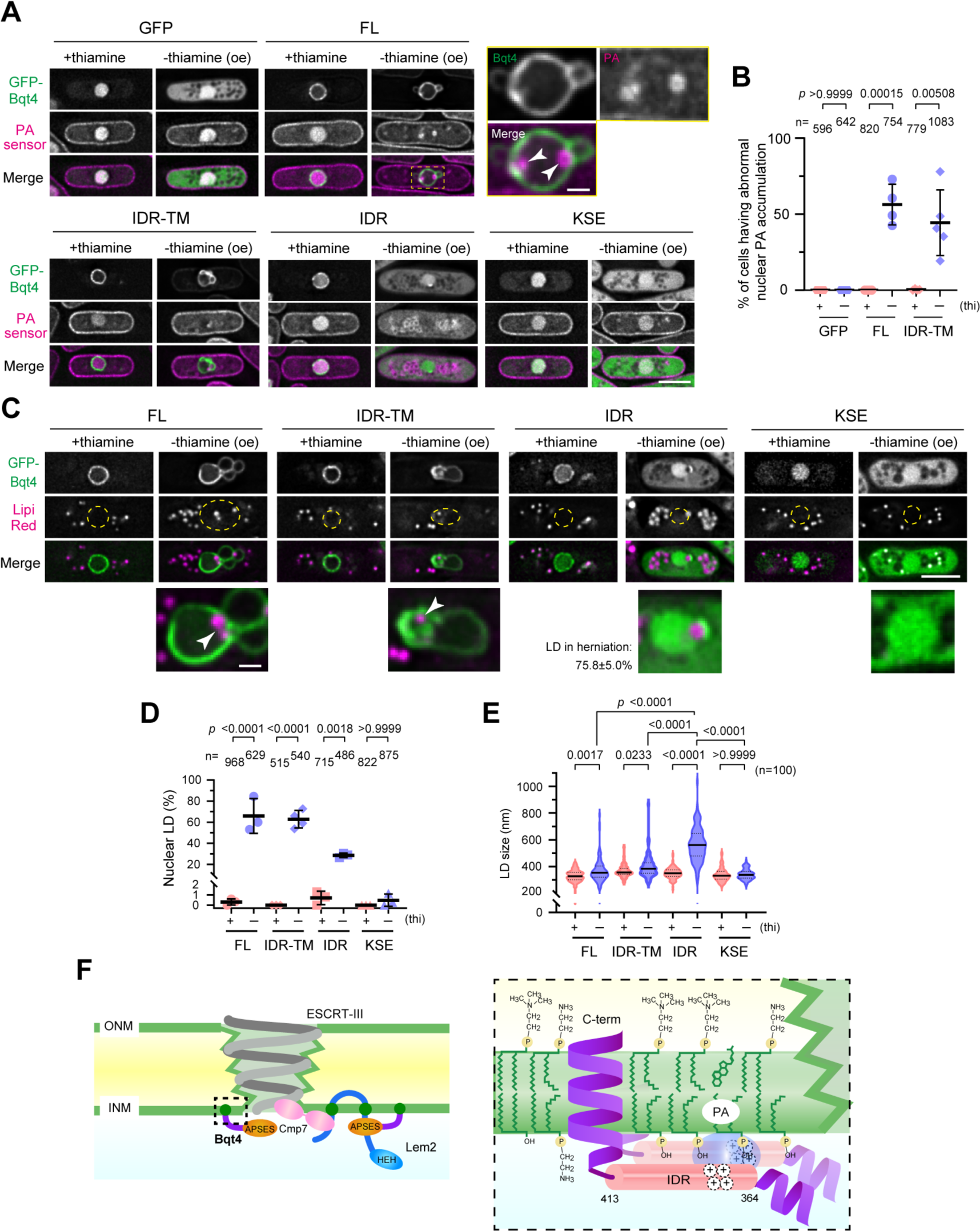
Overexpression of IDR disturbs intracellular lipid distribution. **A.** Visualization of PA in cells overexpressing Bqt4 fragments. The mCherry-tagged PA sensor based on *S. cerevisiae* Opi1 was expressed in *S. pombe* cells. The Bqt4 fragments indicated on the top were overexpressed by culturing the cells in EMMG medium in the presence or absence of thiamine (labeled “+thiamine” and “-thiamine (oe)”, respectively). The scale bar represents 5 μm. The dashed yellow square region in FL is enlarged at the right. Arrowheads indicate the PA-enriched region at the NE. The scale bar for the enraged images represents 1 μm. **B.** Quantification of the cells containing nuclear phosphatidic acid (PA). Percentage of the cells containing a nuclear PA in (A) was determined from three to five independent experiments. The horizontal bold lines and whiskers represent the mean and standard deviation, respectively. The statistical significance was assessed using unpaired two-tailed Student’s *t*-test. Total cell numbers counted (n) are shown on the top. **C.** Intracellular distribution of lipid droplet (LD). Cells overexpressing the Bqt4 fragments were stained with an LD stainer, LipiRed, and the subcellular localization was observed using fluorescence microscopy. Yellow circles indicate nuclear position. Arrowheads indicate nuclear LD. The scale bar represents 5 μm. The bottom row images are enlarged images around the nucleus. The scale bar for the enlarged images represents 1 μm. **D.** Quantification of the cells containing a nuclear LD. Percentage of the cells containing a nuclear LD in (C) was determined from three or four independent experiments. The horizontal bold lines and whiskers represent the mean and standard deviation, respectively. The statistical significance was assessed using one-way ANOVA (*F*=62.55, *p*<0.0001), followed by Tukey’s test. Total cell numbers counted (n) are shown on the top. **E.** Quantification of LD size. The size of LDs observed in (**b**) was measured as follows: measure the line profile of each LD, fit the profile as Gaussian, and then calculate the full width of half maximum (FWHM) of the LD. The FWHM of 100 LDs from at least five cells was measured and plotted as a violin plot. The horizontal bold and dotted lines indicate the median and quartiles, respectively. The statistical significance was assessed using one-way ANOVA (*F*=120.9, *p*<0.0001), followed by Tukey’s test. **F.** Working model. Bqt4 potentially supports NE sealing via two distinct pathways by accumulating PA to the INM. First, Bqt4 generates a preferable micro membrane environment for robust ESCRT-III loading. Bqt4 can enrich PA and Lem2, both of which facilitate and stabilize Cmp7 loading to the NE rupture sites. Second, Bqt4 supplies phospholipids by controlling the balance of lipid synthesis in the INM. Hence, Bqt4 has a broad role in NE maintenance.

As NE proteins are involved in regulating the balance between phospholipid synthesis and energy storage by generating triacylglycerol (TAG) (Romanauska and Köhler, 2018), our results suggest that Bqt4 accumulation forms LDs in the nucleus. Thus, we examined the presence or absence of nuclear LDs in Bqt4-overexpressing cells as a counterpart for phospholipid synthesis. As expected, the number of cells with nuclear LDs increased in the cells overexpressing Bqt4-FL (Fig. 6C and D; 65.9%; see arrowhead in FL) compared to the control cells under the suppressive conditions (0 – 0.3%). Overexpression of the IDR-TM and IDR region also led to an increased number of cells with nuclear LDs (Fig. 6C and D; 62.9% and 28.6%, respectively). Most of the nuclear LDs observed in the cells overexpressing IDR were localized in the herniated NE (Fig. 6C; 75.8 ± 5.0%; arrowhead in IDR) that correspond to a vacuole-like structure observed in the CLEM analysis (compare Fig. 3B with 6C). We also noticed that the LD size in cells overexpressing FL, IDR-TM, and especially IDR, was larger than that in cells under suppressive conditions (Fig. 6C and E). These results suggest that Bqt4 accumulation accelerates TAG synthesis in the NE. As both membrane overproliferation and LD formation were accelerated, Bqt4 possibly plays a role in orchestrating the balance of lipid synthesis via PA accumulation in the INM.

## Discussion

In this study, we uncovered the non-telomeric vital function of Bqt4 in the context of the synthetic lethal phenotype of *lem2^+^* and *bqt4^+^* double deletions. Bqt4 was initially found to anchor telomeres to the NE via its association with the telomeric protein Rap1 (Chikashige et al., 2009). Bqt4 plays a vital role in the absence of Lem2 because the double deletion of *bqt4*^+^ and *lem2*^+^ is lethal (Tange et al., 2016). However, the telomeric function of Bqt4 is not vital, because double deletion of *rap1*^+^ and *lem2*^+^ is viable (Tange et al., 2016). We found that Bqt4 plays a vital role in maintaining NE structures by interacting with specific phospholipids such as PA. Excessive accumulation of Bqt4 on the NE caused membrane overproliferation and nuclear LD formation, which are deleterious to cells (Figs. 2 and 6). Our previous reports showed that excess amounts of Bqt4 are degraded via the ubiquitin-proteasome system, whereas Bqt4 bound to Bqt3 is protected from degradation (Chikashige et al., 2009; Le et al., 2023). We speculate that the protein level of Bqt4 is tightly regulated to maintain a functionally relevant lipid composition in the INM.

The properties of Bqt4 identified in this study allow us to explain why NE ruptures occur in cells lacking Lem2 and Bqt4, and how Bqt4 supports the repair of NE ruptures. We propose that Bqt4 plays a role in NE maintenance via two distinct pathways: (1) production of a preferable microenvironment at the NE rupture sites for the robust association of the ESCRT-III complex, which is an executor of NE repair (Olmos et al., 2015; Raab et al., 2016; Vietri et al., 2015), and (2) lipid supply via the collection of the phospholipid precursor PA into the INM (Fig. 6F).

First, Bqt4 recruits Lem2 (Hirano et al., 2018) and PA (this study) to the NE, playing an important role in functional ESCRT-III complex formation and consequently producing a preferable microenvironment for repairing the ruptured NE membranes. Cmp7/Chm7/CHMP7 accumulates at the rupture sites and recruits other ESCRT-III components, such as Vps32 and Vps24. Lem2 acts as a nuclear adaptor to recruit Cmp7 to rupture sites (Gu et al., 2017; Von Appen et al., 2020). A recent study showed that Chm7 directly binds to PA, and that this PA-binding ability is essential for the binding of Chm7 to the NE (Thaller et al., 2021). As PA preferentially accumulates at the NE holes, which have a high membrane curvature because of their small head group, the ESCRT-III complex is efficiently formed at the NE holes. Moreover, Bqt4 interaction with Lem2 and PA in *S. pombe* infers that Bqt4 recruits and accumulates them to the NE rupture sites to assist Cmp7 in NE sealing, thus supporting the arrangement of preferable membrane conditions for the ESCRT-III complex to seal NE ruptures.

Second, Bqt4 supplies lipids as membrane resources during NE sealing and mitosis. As PA is a precursor of phospholipids, PA in INM produces major structural membrane components, PC and PE; thus, the accumulation of PA by Bqt4 can accelerate membrane expansion. Membrane overproliferation caused by Bqt4 accumulation was similar to that observed in Ned1/lipin and CTDNEP1/CENP-1/Nem1-Spo7 mutants (Fig. 2 and Supplementary Movie 1) (Bahmanyar et al., 2014; Penfield et al., 2020; Santos-Rosa et al., 2005; Tange et al., 2002), supporting this idea. In fungi, PA is utilized for cytidine diphosphate diacylglycerol (CDP-DAG) synthesis by cytidylyltransferase Cds1 (Bbl1 in *S. pombe*), which is a major upstream reaction in PC and PE synthesis (Han et al., 2006). The Ned1 and Nem1-Spo7 complexes are involved in the conversion of PA to DAG in the lipid storage pathway, which antagonizes CDP-DAG synthesis in terms of PA utilization (reviewed in (Carman and Han, 2009)). Loss-of-function mutations in either Ned1 or Nem1-Spo7 have been reported to cause abnormal nuclear membrane expansion (Santos-Rosa et al., 2005; Tange et al., 2002). In the present study, we observed membrane overproliferation (Fig. 2), PA accumulation (Fig. 6A), and LD formation in the nucleus (Fig. 6B) upon the accumulation of Bqt4 in the NE. These results suggest that Bqt4 activates both membrane expansion and lipid storage pathways by retaining PA in the INM that diffuses from the ER. When the NE is ruptured, membrane resources are required to fill the holes, and Bqt4 supports the phospholipid supply. In addition, as Bqt4 interacts with enzymes involved in lipid metabolism in cooperation with Lem2 (Hirano et al., 2023a), Bqt4 can also supply various membrane resources to NE holes. Considering that NE sealing mechanisms can repair micron-scale holes larger than the ESCRT-III spiral filament diameter (approximately 30-50 nm) (McCullough et al., 2015; Penfield et al., 2020), Bqt4 likely supports the repair of NE holes of various sizes by mediating membrane expansion.

Owing to the two Bqt4-mediated membrane repair processes discussed above, the property of IDR to form molecular clusters with PA appears to be advantageous for gathering the necessary components for NE sealing. We have demonstrated that the electrostatic interaction of the KSE region with PA enhances cluster formation via the hydrophobic interaction of IDR-C *in vitro* (Fig. 5). This cluster-forming ability of Bqt4 may play a crucial role in NE maintenance, as supported by the observations in cells carrying mutant Bqt4 proteins. These mutants, characterized by a reduced positive electrostatic potential of the KSE region, show a loss of viability associated with NE ruptures (Figs. 1G, 1H, 4C and Supplementary Fig. 7C). The significance of molecular clustering, including phase separation, of INM proteins for NE maintenance has also been reported in other species: Lem2 undergoes phase separation via its low-complexity domain, which is crucial for NE maintenance in human cells (Von Appen et al., 2020). Considering this, the efficient gathering of essential components via molecular clustering of INM proteins appears to be a conserved mechanism for NE maintenance across various organisms. Bqt4 was the first identified INM protein to regulate INM properties via molecular clustering with lipids. Although Bqt4 is specific to the *Schizosaccharomyces* genus, similar membrane-bound proteins with disordered regions that interact with lipids may serve functional homologs in other eukaryotes.

## Methods

### Yeast strains and culture media

All *S. pombe* and *S. cerevisiae* strains used in this study are listed in Supplementary Table S1. Edinburgh minimal medium with glutamate (EMMG) or EMMG5S (EMMG with 0.225 mg/mL adenine, leucine, uracil, lysine, and histidine) was used as the minimum medium, and the rich medium was yeast extract with supplements (YES) (Moreno et al., 1991). Cells were cultured in an appropriate medium at 30°C for 3-6 days, depending on their growth. Cell strains carrying *lem2*Δ were maintained in EMMG to avoid genomic instability, which occurs in rich medium, as previously reported (Hirano et al., 2018; Hirano et al., 2020; Kinugasa et al., 2019; Tange et al., 2016). For selection, 100 μg/mL G418 disulfate (Nacalai Tesque, Kyoto, Japan), 200 μg/mL hygromycin B (FUJIFILM Wako Pure Chemical Corp., Osaka, Japan), 100 μg/mL nourseothricin sulfate (WERNER BioAgents, Jena, Germany), and 100 μg/mL blastcidin S (FUJIFILM Wako Pure Chemical Corp.) were added to the media.

### Gene disruption, integration, and tagging

Gene disruption, integration, and tagging were performed using a two-step PCR method for direct chromosome integration as previously described (Bähler et al., 1998; Wach, 1996). Briefly, for the first-round PCR, ∼500 bp genomic sequences upstream and downstream of the open reading frames (ORFs) of interest were amplified through PCR using KOD One (TOYOBO, Osaka, Japan; Cat. #KMM-201). These PCR products were used as primers for the second round of PCR to amplify a template sequence containing the selection markers. The resulting PCR products were transformed into *S. pombe* cells for disruption, integration, and tagging, and transformants were selected on an appropriate selection plate. The obtained strains were confirmed for correct constructions of disruption, integration, and tagging via genomic PCR using KOD Fx Neo (TOYOBO; Cat. #KFX-201) at both the 5′ and 3′ ends. In addition, we performed genomic PCR on the ORF of the gene of interest to confirm the absence of the ORF in the genome.

### Plasmid construction

All plasmids used in this study were constructed using the NEBulider (New England Biolabs, Ipswich, USA; Cat #E2621L) according to the manufacturer’s protocol. Restriction enzymes were purchased from TaKaRa Bio Inc. (Kusatsu, Japan) and New England Biolabs (Ipswich, MA, USA). cDNA sequences encoding Bqt4 fragments were amplified via PCR from the pCSS18 plasmid (Chikashige et al., 2009) and cloned into the authentic *bqt4* promoter-harboring pCSS41 at *Bam*HI site. Deletion and point mutants of Bqt4 were generated using inverse PCR. After confirming the DNA sequences, these mutants were cloned into *nmt1* promoter harboring pCST3 at *Bam*HI site or pGADT7-AD (Clontech) at *Eco*RI site. cDNA sequence of Bqt3 was amplified and cloned into pGBKT7-DB (Clontech) at *Eco*RI and *Bam*HI sites for the yeast two-hybrid assay. To express the Bqt4 protein in *E. coli*, cDNA sequences encoding GFP, GFP-IDR (364-413 aa), GFP-KSE (364-383 aa), and GFP-IDR-C (384-413 aa) were amplified via PCR and cloned into the pColdI vector (Clontech) at *Bam*HI site.

The DNA sequence of the PA biosensor *S. cerevisiae* (NLS) Opi1^111-189^2xAH^G120w^ (Foo et al., 2023), was synthesized using GenScript (Piscataway, NJ, USA) and cloned into the pYK172 vector at *Bam*HI site.

### Spot assay

*S. pombe* cells were cultured overnight in EMMG at 30°C to attain the logarithmic growth phase. Five-fold serially diluted cells were spotted on EMMG plates supplemented with or without 10 μM thiamine (initial cell numbers spotted are 4.0×10^3^ cells) and cultured at 30°C for 3-4 days.

### Yeast two-hybrid assay

The yeast two-hybrid assay was performed according to the manufacturer’s protocol (Clontech MATCHMAKER System Manual, Clontech). Plasmids harboring Bqt4 or Bqt3 were co-transfected into *Saccharomyces cerevisiae* AH109 strain, and the cells were grown on synthetic defined (SD) media with dropout supplements -Leu/-Trp (cat #630417). To examine protein interactions, suspensions of the transformants (1.0×10^6^ cells) were spotted onto SD plate media with dropout supplements, -Leu/-Trp or -Ade/- His/-Leu/-Trp (cat #630428) (Takara Bio), and incubated at 30°C for 3 days.

### Protein expression and purification

Plasmids harboring His-GFP, His-GFP-IDR, His-GFP-KSE, His-GFP-IDR-C were introduced into the BL21 (DE3) pLysS strain. The cells were cultured in LB medium at 37 °C until OD_600nm_ ∼0.5. After adding the final 0.1 mM Isopropyl β-D-1-thiogalactopyranoside, the protein expression was induced at 16 °C for 24 h. The cells were harvested, washed, and then resuspended in resuspension buffer (phosphate-buffered saline [PBS] containing 0.1% Triton-X100 and 1 mM phenylmethylsulfonyl fluoride). After sonication and centrifugation, the supernatant supplemented with final 10 mM imidazole was incubated with His-Select Nickel Affinity Gel (P6611, Lot No. 110M5166, SIGMA) for 1.5 h at 4 °C with rotation. The gel was washed three times with resuspension buffer containing 10 mM imidazole, and the bound proteins were eluted by stepwise treatment with 50, 100, and 300 mM imidazole. The purity and quantity of the eluted proteins were assessed through SDS-PAGE followed by CBB staining with BSA as a standard.

### Lipid binding assay

Membrane lipid strip (P-6002) was purchased from Echelon Biosciences (Salt Lake City, UT, USA). The membrane was incubated with blocking buffer (PBS containing 0.1% Triton X-100 and 3% fatty-acid- and globulin-free BSA [A7030, Lot No. SLCH7959, SIGMA]) for 1 h at room temperature (approximately 26-28°C; RT), and then incubated with 1 µM of His-GFP-IDR, or His-GFP in the blocking buffer for 1.5 h at RT. After washing three times with washing buffer (PBS containing 0.1% Triton X-100), the fluorescence emitted from bound proteins was detected by using ChemiDoc Touch (Bio-Rad) equipped with a filter set for Alexa488.

For the liposome-binding assay, POPA (1-palmitoyl-2-oleoyl-*sn*-glycero-3-phosphate; Cat. #840857C), POPC (1-oleoyl-2-palmitoyl-*sn*-glycero-3-phosphocholine; Cat. #850475C) and POPE (1-palmitoyl-2-oleoyl-sn-glycero-3-phosphoethanolamine; Cat. #850757C) in chloroform solution were purchased from Avanti Polar Lipids (NJ, USA). Lipid solutions were mixed at specific ratios in round-bottomed glass tubes. The solutions were blown dry under Argon gas in heating blocks at 45°C and resulting lipid film was further dried under vacuum overnight. HEPES buffer (10 mM 4-(2-hydroxyethyl)-1-piperazineethanesulfonic acid (HEPES)-NaOH (pH 7.6)) was added to the tubes and hydrated with 10 mM liposome suspensions. The suspensions were incubated in a 55°C water bath for 10 min and then shaken with a vortex mixer for 30 s. This procedure was repeated three times. The freeze-thaw cycles were repeated at least 10 times between liquid nitrogen and a 55 °C water bath. The liposome solutions were transferred to 5 × 41 mm ultracentrifuge tubes (Beckman Coulter, CA, USA; Cat. #344090). Two millimolar liposomes were mixed with the indicated concentration of proteins to a total volume of 150 µL. The mixed solutions were incubated at 30°C for 1 h, and then gently mixed an equal volume of 80% Histodentz (Sigma-Aldrich, MO, USA; Cat. #D2158) in HEPES buffer and adjusted to 40% Histodenz. The mixture was overlaid first with 250 µL of 30% Histodentz in HEPES buffer, then with 50 µL HEPES buffer. Samples were spun at 200,000 × g at 4°C for 4 h in an MLS-50 rotor (Beckman Coulter), and 60 µL of top layer was collected. After adding 18 µL of 4 × Laemmli sample buffer and 2 µL of 2-mercaptoethanol to the collected fraction, samples were denatured at 95°C for 5 min. Samples were electrophoresed in a 10% gel (Bio Craft, Tokyo, Japan; Cat. #SDG-521) and visualized using 2D-Silver Stain Reagent II (Cosmo Bio, Tokyo, Japan; Cat. #423413).

### Determination of Bqt4 concentration in the NE

Cells expressing GFP-Bqt4 under the *bqt4* or *nmt1* promoter were attached to a glass-bottom dish (MatTek, Cat #P35G-1.5-14-C) coated with soybean lectin (Sigma-Aldrich, St. Louis, USA; Cat #L1395). Serially diluted purified His-GFP was dropped onto the cells, and then the cells and GFP were simultaneously observed through spinning disk confocal microscopy as described in “Microscopic observation.” The fluorescence intensities outside the cells were quantified, plotted, and used as a standard to quantify the Bqt4 concentration.

### In vitro condensation assay

For condensation assay with PA, the purified GFP, GFP-IDR, GFP-KSE, and GFP-IDR-C proteins (final 2 μM) were mixed with 0, 20, 50, 100 nmol PA or PC on ice, and 2 μL of the mixture was put onto a cover glass (No. 1-S, Matsunami, Osaka, Japan) immediately. For condensation assay in crowding condition, the GFP-Bqt4 fragments (final 2 or 8 μM) were diluted into crowding buffer (25 mM Tris-HCl pH 7.5, 5 mM MgCl_2_, 1 mM dithiothreitol, 5% glycerol) with or without 5% PEG8000. Final 0.5 M NaCl, 2 M urea, 0.2 M arginine, 10% 1,6-HD, 0.1% SDS, and 0.1% triton-X100 were added to condensation buffer with PEG8000 to prevent aggregate formation. The formed aggregates were observed through spinning disk confocal microscopy as described in “Microscopic observation.”

### Microscopic observation

*S. pombe* cells were observed using a DeltaVision Elite system (GE Healthcare Inc., Chicago, USA) equipped with a pco.edge 4.2 sCMOS (PCO, Kelheim, Germany) and a 60× PlanApo N OSC oil-immersion objective lens (numerical aperture [NA] = 1.4, Olympus, Tokyo, Japan). Intracellular localization of GFP-S65T (designated “GFP” throughout this study) and mCherry fusion proteins was observed in living cells. For time-lapse imaging, cells were cultured overnight in EMMG medium at 30°C to attain the logarithmic growth phase before being placed in a glass-bottom culture dish. The cells were attached to the glass via soybean lectin and covered with EMMG medium.

For the condensation assay and quantification of the GFP-Bqt4 concentration in NE, a spinning disk confocal microscope system (CSU-W1; YOKOGAWA, Kanazawa, Japan) equipped with an Orca-Fusion BT sCMOS camera (HAMAMATSU Photonics, Hamamatsu, Japan) and a 60× PlanApo VC oil-immersion objective lens (numerical aperture [NA] = 1.4; Nikon, Tokyo, Japan) was used. The microscope was operated using the built-in NIS-Elements software (v5.42.01).

Chromatic aberration, except for time-lapse observations for multicolor images, was corrected using Chromagnon software (v0.90) with a four-color fluorescence bead (Thermo Fisher Scientific) image as a reference (Matsuda et al., 2018). Optical section images captured by the DeltaVision Elite system were presented after deconvolution using the built-in SoftWoRx software (v7.0.0). The brightness of the images was changed using Fiji (Schindelin et al., 2012) or Adobe Photoshop 2022 software for better visualization and presentation without changing the gamma settings.

FRAP analysis was performed as described previously with slight modifications (Hirano et al., 2018). *S. pombe* cells were attached to a CELLVIEW cell culture dish (Griner Bio-one, Austria; Cat. #627871) coated with soybean lectin. The mobility of wild-type Bqt4 and its mutants was analyzed using an LSM780 confocal microscope system (Carl Zeiss, Jena, Germany) equipped with a 63× oil-immersion objective lens (NA = 1.4). For the FRAP analysis in cells, three images were collected before bleaching (approximately 1% transmission of a 488-nm argon laser, 970 ms/frame, 1.58 μs/pixel, average 2, 512 × 256 pixels, 48 μm pinhole corresponding to 1 airy unit, 7× zoom); the NE signal was bleached using 100% of a 488-nm laser with three iterations, followed by the capture of a further 47 images, using the original setting. For FRAP assay of GFP-IDR aggregate, the following condition was used: a 40× C-Apochromat water-immersion objective lens (NA = 1.2) was used. Five images were collected before bleaching (approximately 0.02% transmission of a 488-nm argon laser, 968 ms/frame, 3.15 μs/pixel, average 1, 512 × 256 pixels, 33.8 μm pinhole corresponding to 1 airy unit, 4× zoom); the part of the aggregate was bleached using 100% of a 488-nm laser with three iterations, followed by the capture of a further 95 images, using the original setting. The fluorescence intensity in the bleached region was quantified using Zen 2.3 SP1 software. Photobleaching during imaging was monitored and normalized before the recovery curve was plotted. The curve was plotted as a relative value and the average fluorescence intensity before bleaching was 1.

FCS measurements were performed using an LSM510META confocor2 (Carl Zeiss) equipped with a 40× C-apochromatic water immersion objective lens. The measurements were performed using the built-in AIM software (version 3.2 SP2). GFP proteins were excited using argon laser (488 nm) and the fluorescence passed a pinhole (70 μm in diameter). The band-pass filter (530-600 nm) was detected using an avalanche photodiode. The fluctuation in fluorescence intensity was measured for 10 s in 10 cycles. The averaged autocorrelation function curves are presented.

### CLEM imaging

CLEM imaging was performed as described previously (Asakawa et al., 2014; Asakawa et al., 2016). Briefly, cells expressing Bqt4 fragments were cultured overnight in EMMG to express Bqt4 proteins. The cells were attached to a dish with a gridded coverslip (Ibidi, Martinsried, Germany) as described above. After live-cell observations, the medium was replaced with a fixative solution (4% paraformaldehyde, 250 mM Hepes-NaOH, pH 7.2) for 10 min at RT (approximately 26-28°C). The cells were monitored via fluorescence microscopy using the DeltaVision system before and during fixation to confirm that the structures observed before fixation were preserved after fixation. The cells were further fixed overnight with fresh fixative (2% glutaraldehyde in 0.1 M sodium phosphate buffer, pH 7.2) at 4°C. After washing with phosphate buffer, the cells were post-fixed in 1.2% KMnO_4_ overnight, dehydrated, and embedded in epoxy resin. Serial sections of 80 nm thickness were stained with uranyl acetate and lead citrate and analyzed using a transmission electron microscope (JEM1400Plus, JEOL, Japan) at 100 kV.

### Western blotting

Cells cultured in the appropriate minimum medium, as described above, were harvested at the mid-log phase using centrifugation. The cells were lysed in 1.85 N NaOH for 15 min on ice. Proteins were precipitated by adding 27.5% trichloroacetic acid. After washing the precipitated proteins twice with ice-cold acetone, they were resolved in 2× Laemmli SDS sample buffer without dye. The protein concentration was quantified using a bicinchoninic acid assay (Thermo Fisher Scientific, Waltham, MA, USA; cat. #23225) according to the manufacturer’s protocol. The same amount of total protein (10 μg) was subjected to SDS-PAGE and then transferred to a polyvinylidene fluoride membrane. The fluorescent proteins were probed with anti-GFP monoclonal (JL8, 1:2,000 dilution; TaKaRa Bio Inc.) and anti-RFP polyclonal (PM005, 1:5,000 dilution; MBL, Nagoya, Japan) antibodies, and detected using chemiluminescence (ImmunoStar LD, FUJIFILM Wako Pure Chemical Corp.). The membrane was stained with Amido black to determine the amount of loaded proteins.

### Quantification, statistics, and reproducibility

For phenotypic quantification of NE deformation, images were collected from at least three independent experiments. Percentages of the phenotype were calculated in each experiment (total number of cells examined was> 500), and the values are represented as mean with standard deviation. LD size was measured using default functions in the Fiji software. The fluorescence intensity line profile of each LDs was measured and fitted as a Gaussian, and the full-width at half-maximum of the LD was calculated from the fitted parameter. The values obtained from 100 LDs were plotted as violin plots. Significance was assessed using the chi-square test, Tukey’s test, or unpaired two-tailed Student’s *t-*test, as noted in the figure legends, using GraphPad Prism 9.4.1 or later version.

## Data availability

The data supporting the findings of this study are available in the paper and Supplementary Information files. Additional data sets are available from the corresponding author upon request.

## Acknowledgments

We thank Chizuru Ohtsuki for her technical assistance. This study was supported by JSPS KAKENHI Grant Numbers JP19K06489 and JP20H05891 (to Y. Hirano); JP18K05356 (to K.K.); JP18H05528 (to T.H.), and JP20H00454 and JP23K05636 (to Y. Hiraoka), and also by the grant of CREST of JST (JPMJCR21E6) to TF.

## Declaration of Interests

The authors declare no competing financial or non-financial interests.

## Contact for reagent and resource sharing

Further information and requests for resources and reagents should be directed to Yasuhiro Hirano or Yasushi Hiraoka

---

**Supplementary Fig. S1.**
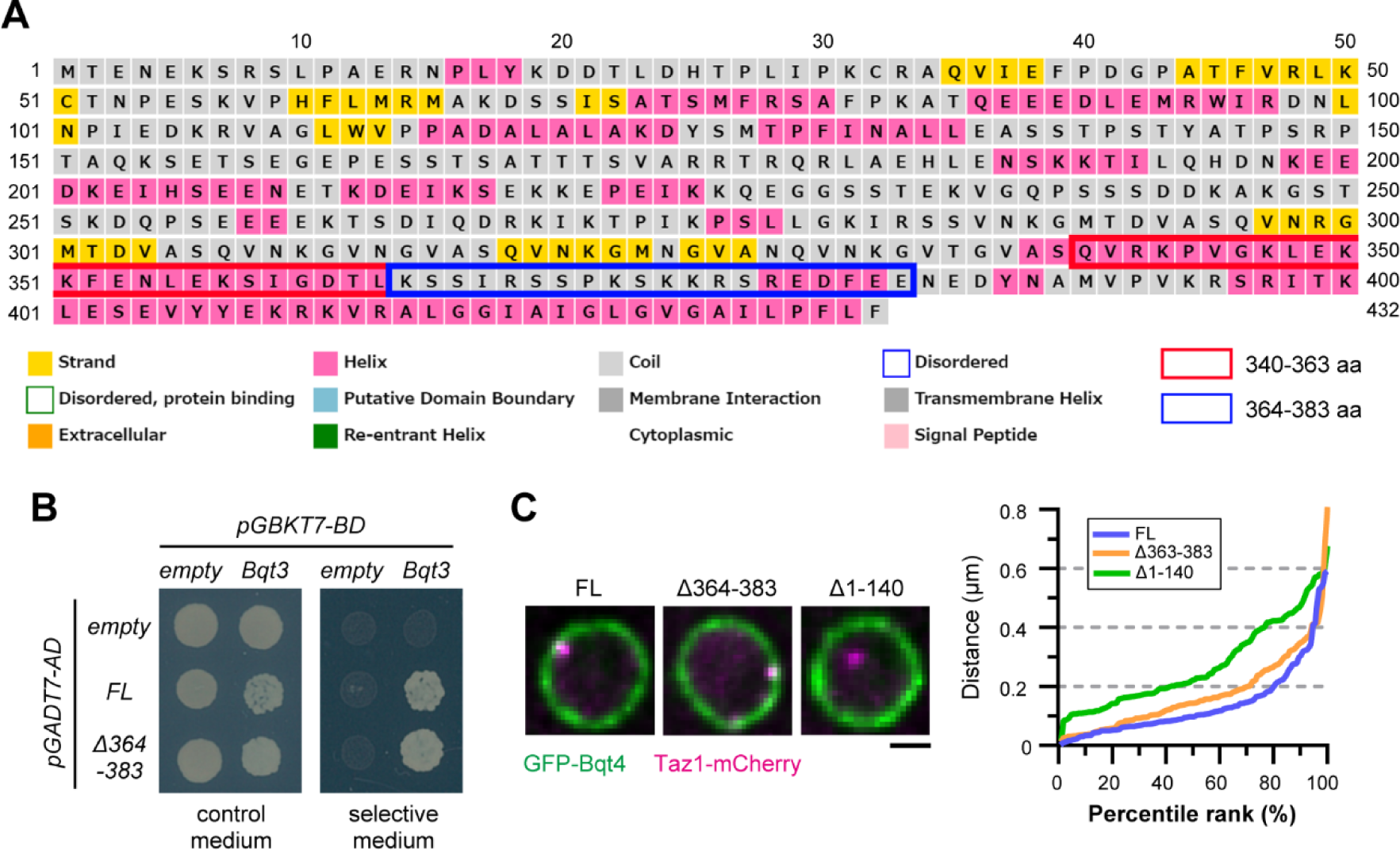
KSE region characteristics. **A.** Secondary structure prediction of Bqt4 by PSIPRED 4.0. The sequences surrounded by red and blue frames indicate 340-363 aa and 364-383 aa, respectively. **B.** Yeast two-hybrid assay. *S. cerevisiae* cells expressing the full-length Bqt4 (FL) and the Bqt4 Δ364-383 fragment with Bqt3 were grown on a medium with different auxotrophy (control and selective medium) for 3 days. Fusion proteins with AD (pGADT7-AD) and BD (pGBKT7-BD) are shown on the left and top, respectively. **C.** Distance between the NE and telomeres. (Left) Full-length (FL) and fragments (Δ364-383 and Δ1-140) of Bqt4 tagged with GFP (green) were co-expressed with Taz1-mCherry as a maker for telomere (magenta). (Right) Distance between the NE to telomeres was measured and plotted as a percentile rank. The scale bar represents 1 μm.

**Supplementary Fig. S2.**
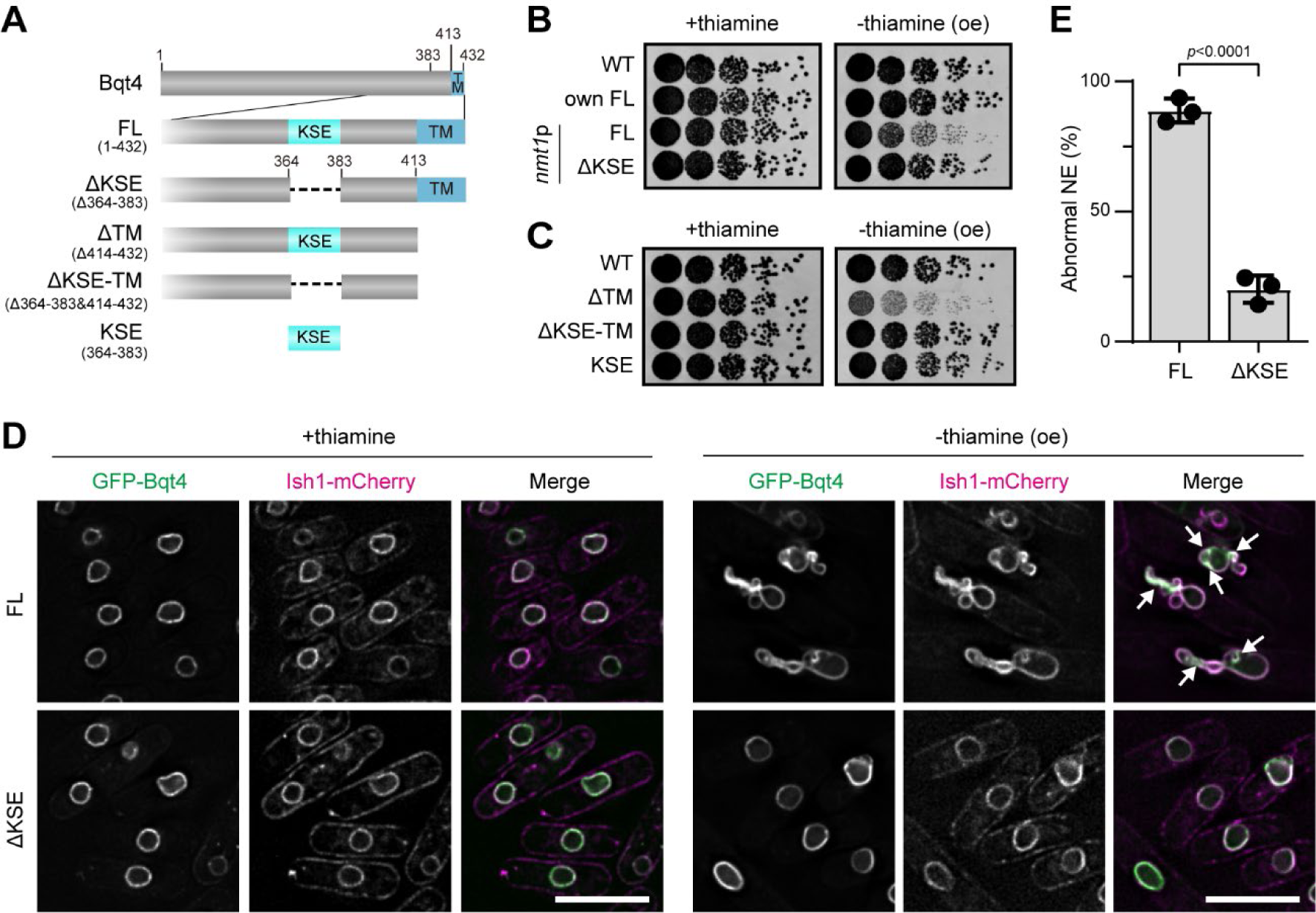
The KSE region is required for cell growth, but insufficient for dominant negative defects. **A.** Schematic diagrams of KSE fragments. TM denotes the transmembrane domain. **B.** Spot assay of the cells overexpressing ΔKSE fragment. GFP-tagged Bqt4 fragments indicated on the left were expressed under authentic (own) or *nmt1* (*nmt1p*) promoters. The growth of the cells was observed as described in Fig. 1A. oe represents overexpression. **C.** Spot assay of cells overexpressing the Bqt4 fragments. GFP-Bqt4 fragments indicated on the left were expressed under the *nmt1* promoter. The growth of the cells was observed as described in (B). **D.** NE deformation by Bqt4 overexpression. GFP-tagged FL or ΔKSE were expressed with Ish1-mCherry as a nuclear membrane marker (Asakawa et al., 2022) in *S. pombe* cells. The cells were cultured in the EMMG medium with or without thiamine (labeled “+thiamine” and “-thiamine (oe),” respectively), and then observed using fluorescence microscopy. Arrows indicate the region of Bqt4 enrichment. The scale bars represent 5 μm. **E.** Quantification of abnormal NE morphology. Percentage of the cells that showed abnormal NE morphology in (D) was quantified from three independent experiments. The mean and standard deviation were shown. The statistical significance was assessed using two-tailed Welch’s *t*-test.

**Supplementary Fig. S3.**
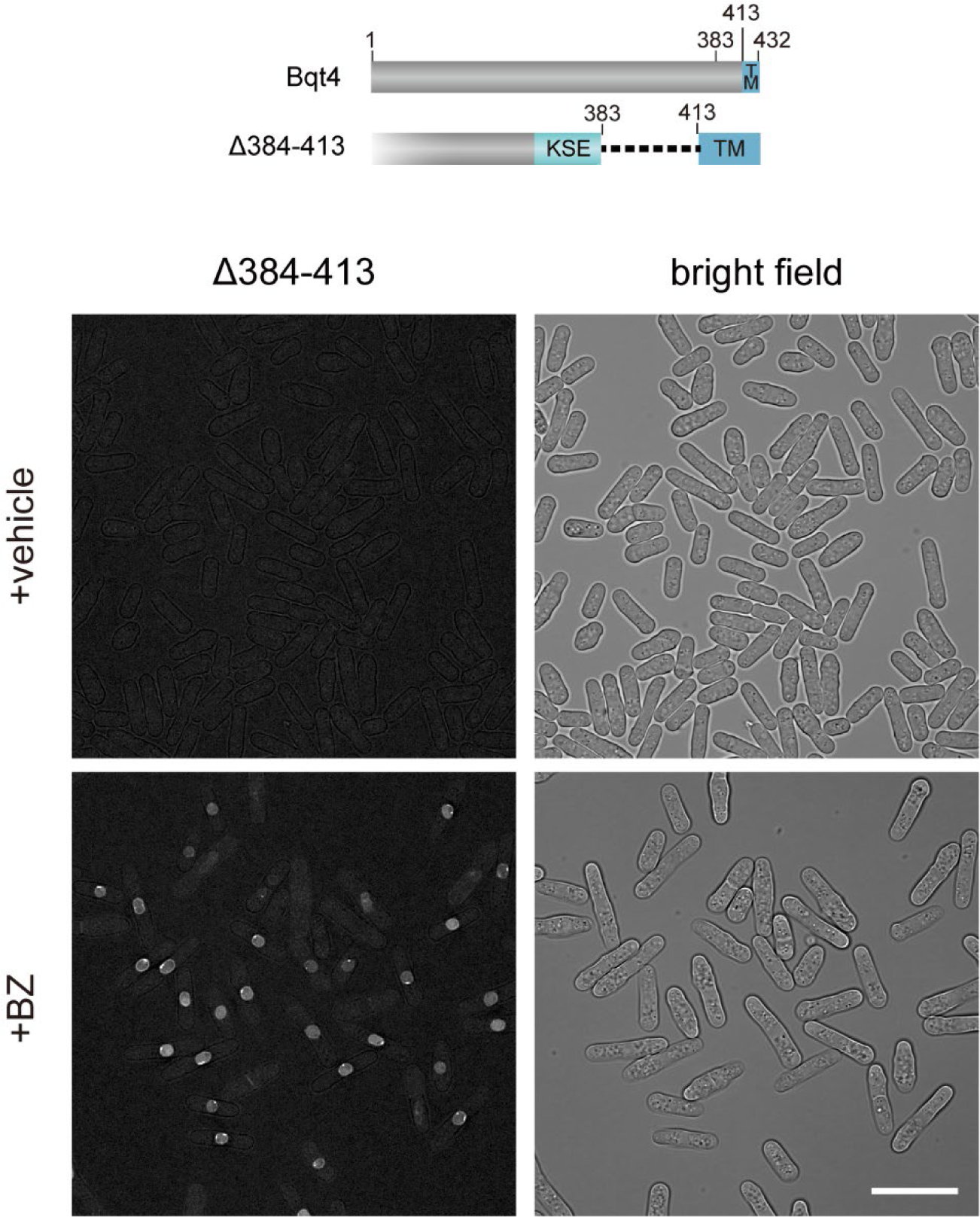
Bqt4Δ384-413 protein fragment undergoes degradation. GFP-tagged Bqt4Δ384-413 fragment was expressed in the *lem2*-shut-off *bqt4*Δ cells and observed in the presence (+BZ) or absence (+vehicle) of 1mM bortezomib, a proteasome inhibitor effective in *Schizosaccharomyces pombe* cells. Bright-field images are shown on the right side. Scale bar represents 20 μm.

**Supplementary Fig. S4.**
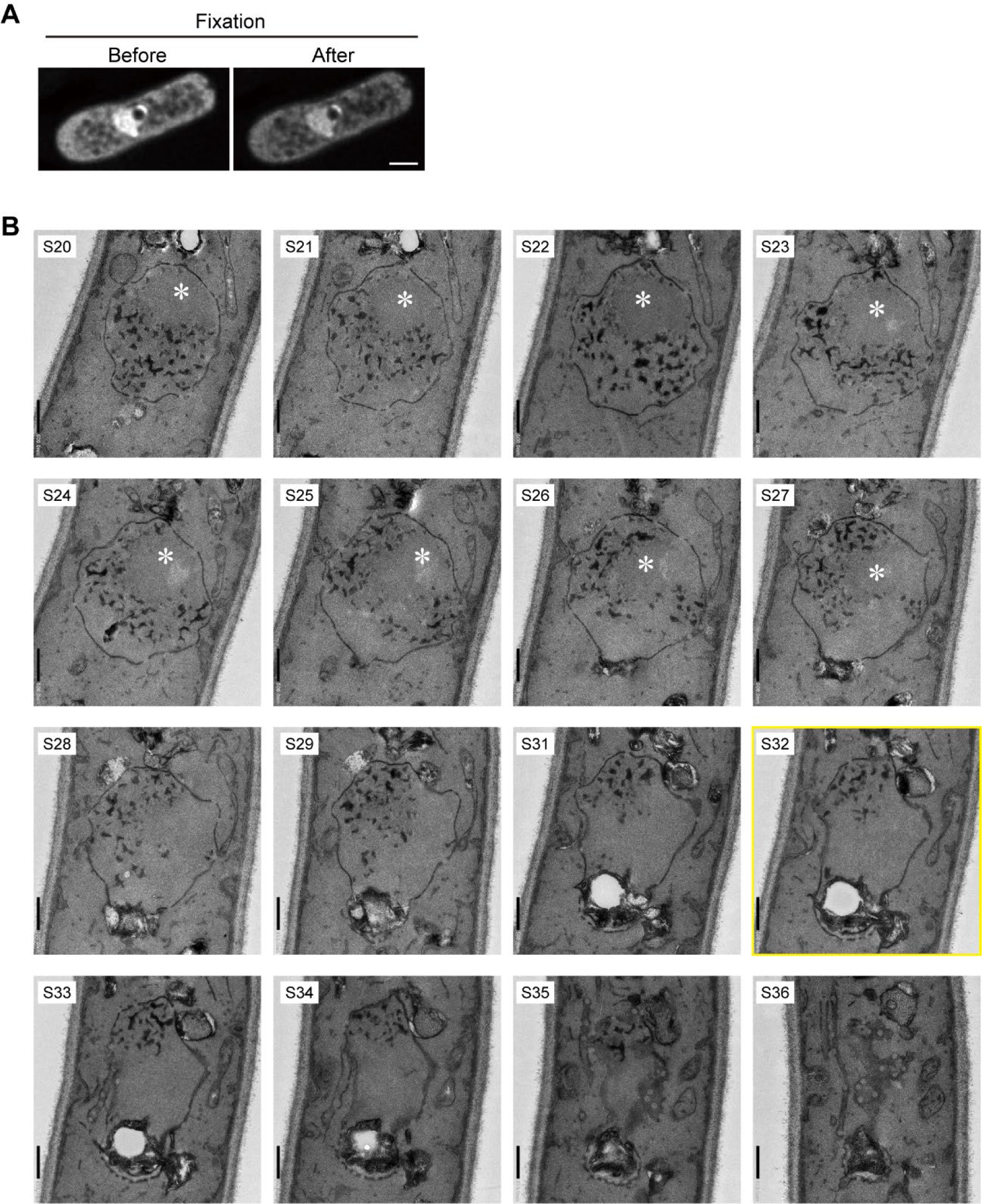
Serial section TEM images of the cell overexpressing IDR. **A.** Fixation did not alter the structure observed in living cells. Cells overexpressing IDR were observed in live (Before) and then fixed with PFA for 10 min. The same region observed in live imaging was observed again after fixation (After). The scale bar represents 2 μm. **B.** Serial-section TEM images of cells overexpressing IDR are shown in Fig. 2H. The numbers on the upper left indicate the section numbers. S32 is shown in Fig. 2H. Asterisk at S20-S27 indicates the nucleolus. The scale bars represent 500 nm.

**Supplementary Fig. S5.**
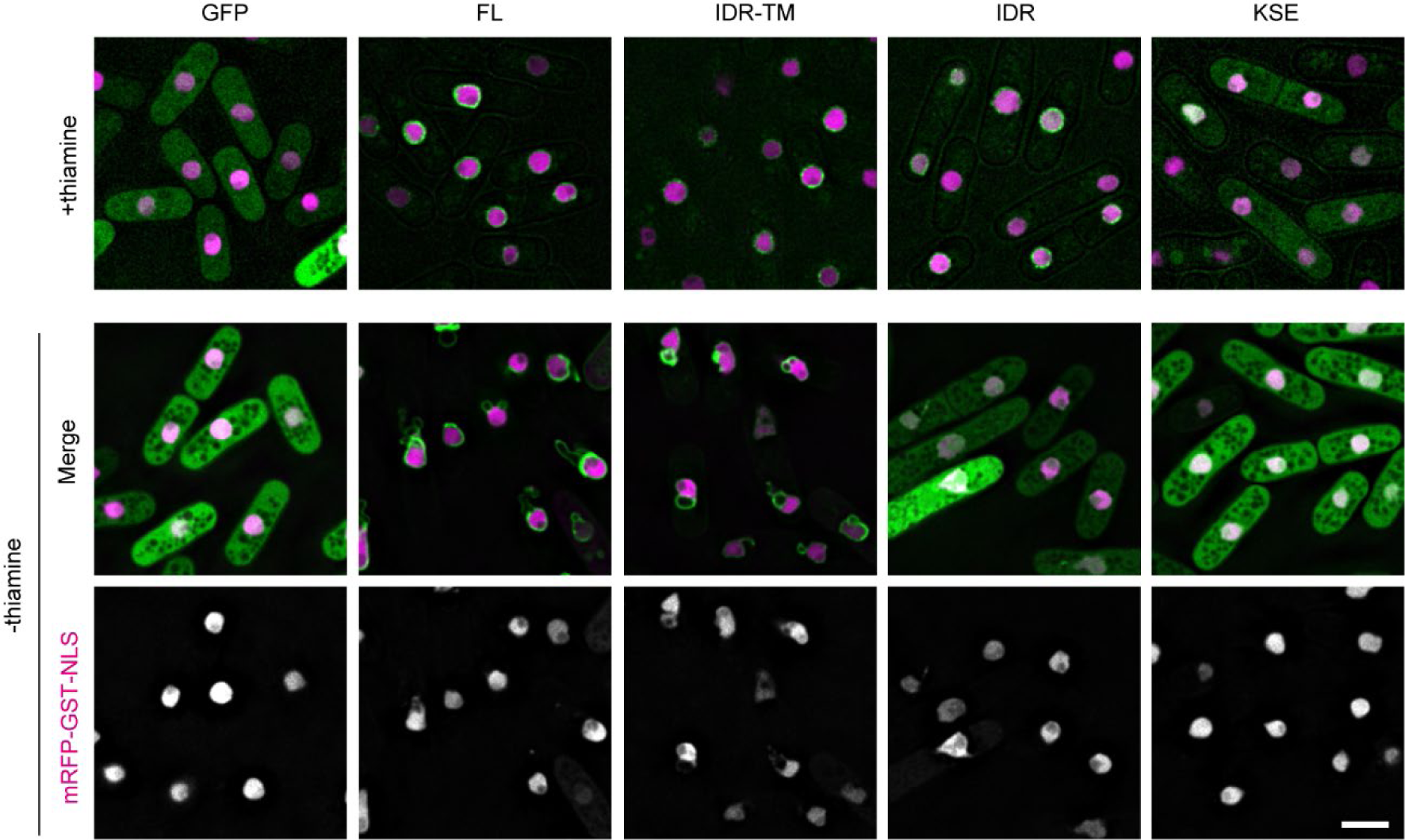
Overexpression of Bqt4 fragments did not disturb the NE integrity. Cells expressing Bqt4 fragments and mRFP-GST-NLS were cultured in the presence or absence of thiamine and observed using fluorescence microscopy. Scale bar represents 5 μm.

**Supplementary Fig. S6.**
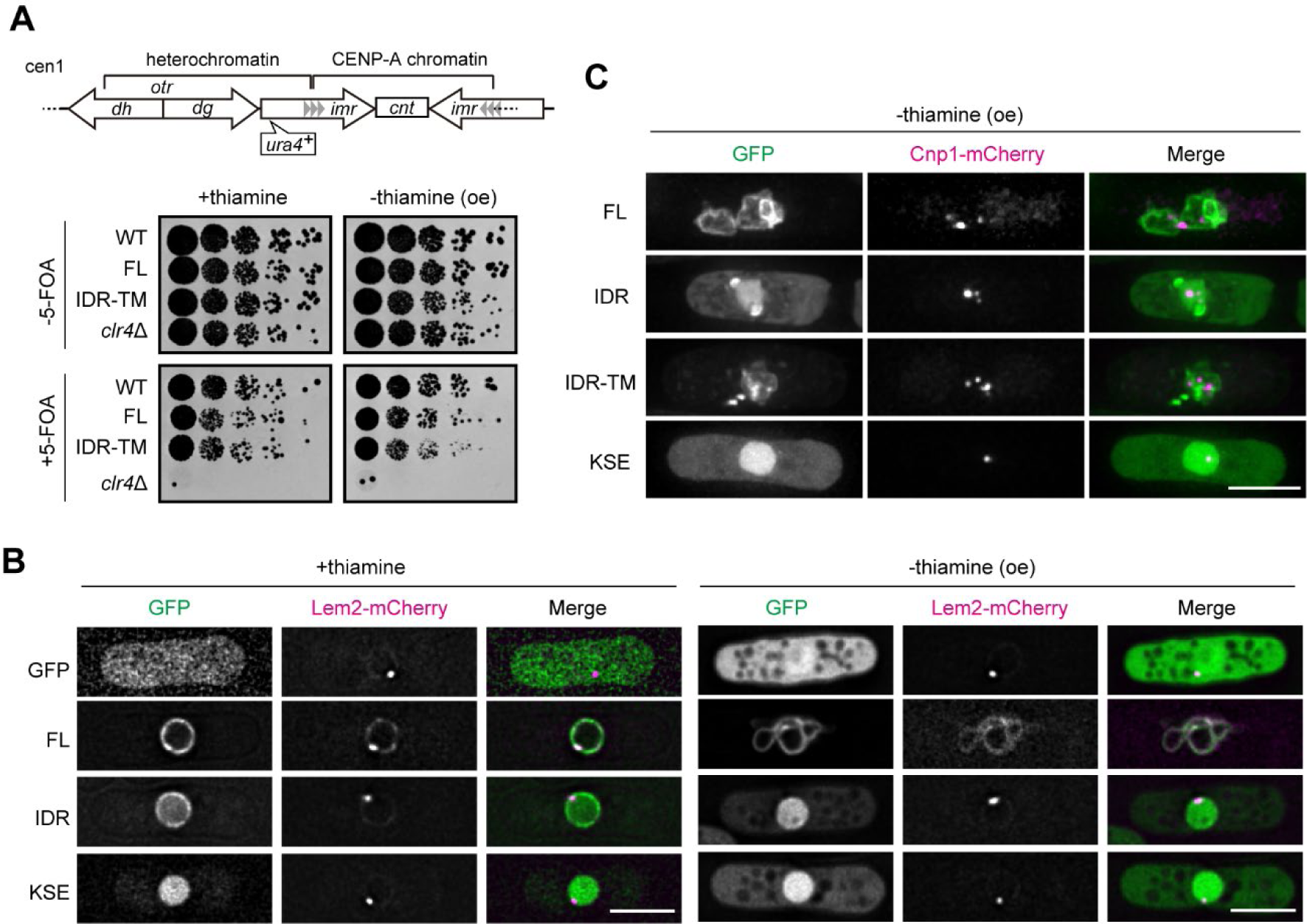
Lem2 did not involve centromeric defects caused by Bqt4 overexpression. **A**. Heterochromatin silencing assay in cells overexpressing GFP-FL or IDR-TM. The cells with increased expression of GFP-FL or -IDR-TM and *clr4*Δ cells were spotted on the EMMG containing 5-FOA with or without thiamine in five-fold serial dilutions and then grown at 30°C for 4 days. **B**. Subcellular localization of Lem2 in the cells overexpressing Bqt4 fragment. GFP-tagged Bqt4 fragments (green) indicated on the left were expressed in the cells expressing mCherry-tagged Lem2 (Lem2-mCherry, magenta). The cells were cultured in the EMMG medium in the presence or absence of thiamine (“+thiamine” and “-thiamine (oe)”, respectively). Lem2-mCherry is accumulated at the centromere in the control cells disrupted for the authentic *bqt4^+^*gene as previously reported (Hirano et al., 2018). The scale bars represent 5 μm. **C.** Distribution of centromeres in *lem2*Δ cells. *lem2*Δ cells expressing the GFP or GFP-Bqt4 fragments indicated on the left (green) and Cnp1-mCherry (magenta) were cultured in EMMG medium and observed using fluorescence microscopy. The scale bar represents 5 μm.

**Supplementary Fig. S7.**
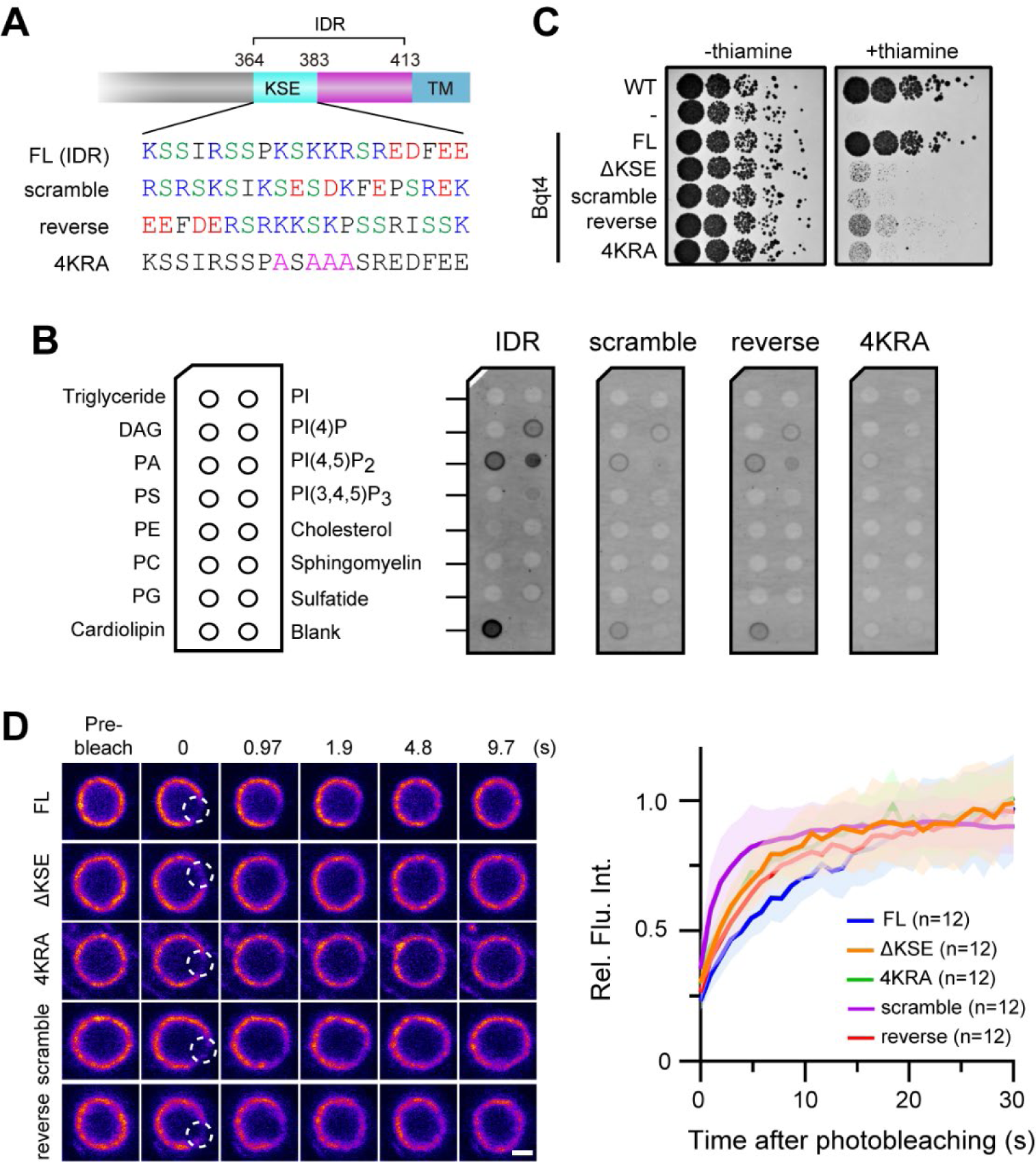
Disturbing the highly positive charge in the KSE region leads to the loss of the Bqt4 function. **A**. Bqt4 proteins used. We constructed Bqt4 mutants with scrambled or reversed amino acid sequences of the KSE region (scramble and reverse, respectively). The 4KRA is the same mutation shown in Fig. 4. These mutations were introduced in full-length Bqt4 (FL) and the IDR fragment (IDR) for spot and FRAP analyses and lipid-binding assay, respectively. Blue, red, green, and pink characters represent positive, negative, polar, and mutated amino acids. **B**. Lipid binding assay. The schematic illustration on the left shows lipids spotted on the membrane lipid strip. His-GFP-tagged IDR, scramble, reverse, or 4KRA was incubated with the membrane and the bound protein was detected using fluorescence. **C**. Spot assay of the mutants in the KSE region. The cells expressing mutants shown on the left were spotted and the growth was observed as described in Fig. 1B. **D**. FRAP analysis of GFP-tagged Bqt4 fragments (FL, ΔKSE, 4KRA, scramble, reverse, and 4KRA). The dashed white open circle region was bleached, and the recovery of fluorescence intensity after bleaching in the circle was measured. Solid lines in the FRAP curve indicate the mean value. The areas on both sides of the solid lines indicate standard deviation. The scale bar represents 1 μm.

**Supplementary Fig. S8.**
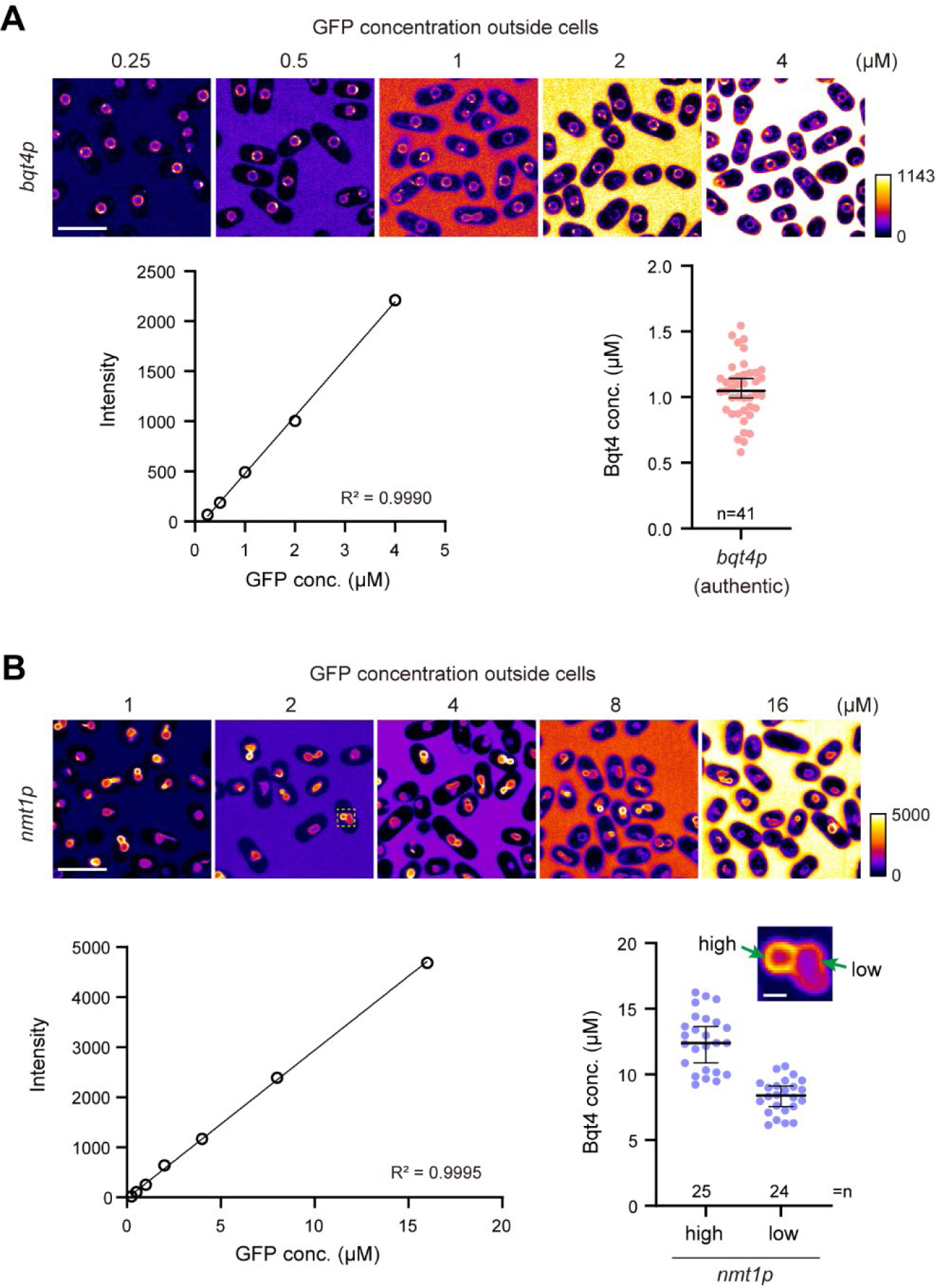
Bqt4 concentration in the NE. Cells expressing GFP-Bqt4 under authentic (A) or *nmt1* (B) promoters were mixed with purified GFP solutions of increasing concentrations indicated on the top and observed using fluorescence microscopy (top panels). A standard curve was constructed by quantifying the intensities outside of the cells (bottom left). Bqt4 concentrations in the NE were determined using a standard curve (bottom right). Since the Bqt4 concentration in the NE expressed under the *nmt1* promoter was not uniform, we categorized it into high- and low-concentration regions (inset of (B)). The horizontal line and whiskers in the plots denote median and 95% confidence intervals. The scale bars represent 10 μm for the top row images in (A) and (B), and 1 μm for the inset of (B).

**Supplementary Fig. S9.**
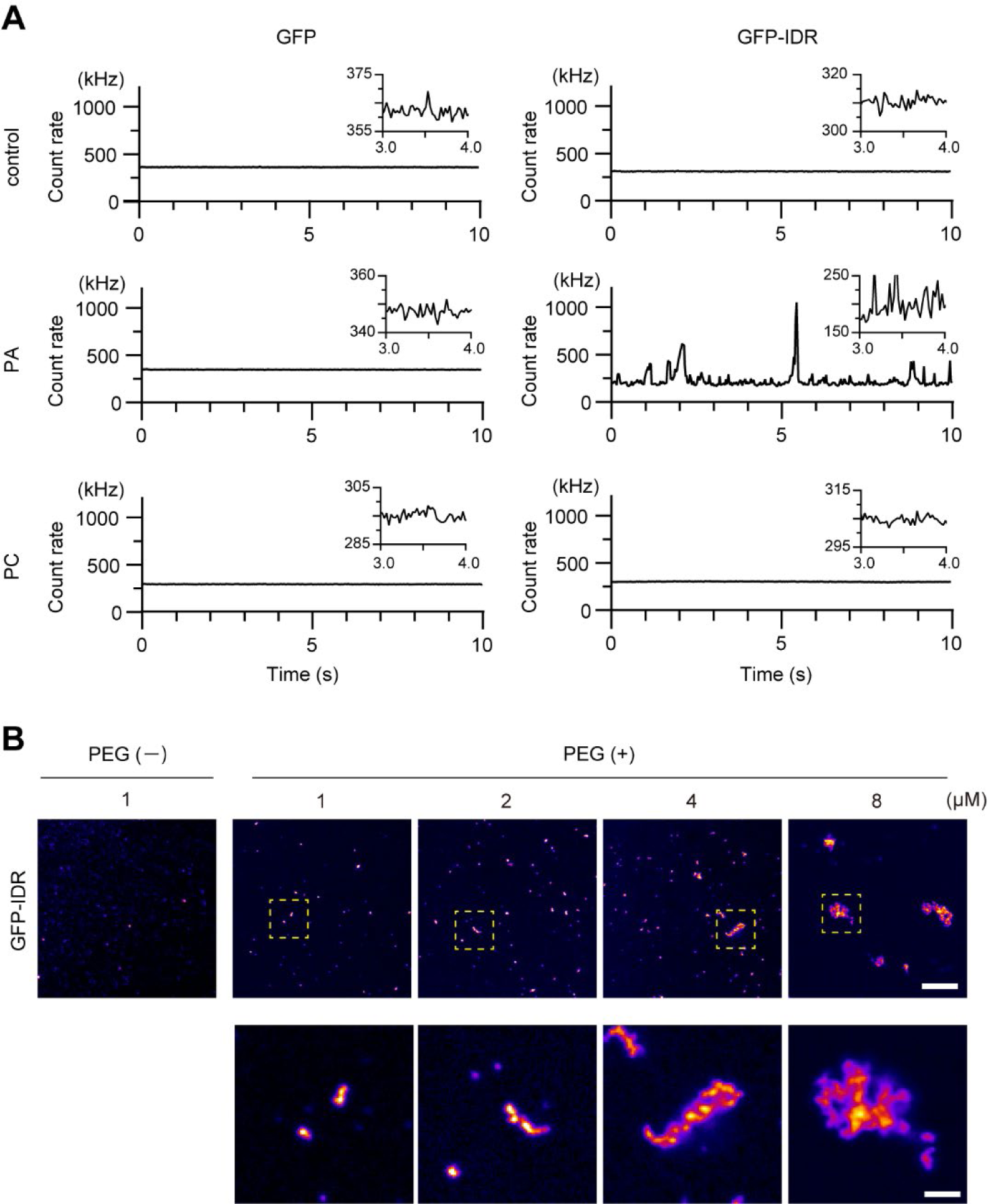
Molecular clustering of GFP-IDR. **A.** Fluorescence intensity fluctuation during FCS measurements. GFP (left) and GFP-IDR (right) were mixed with 100 nmol PA or PC, and then the mixture was analyzed using FCS. Inlets are enlarged images of the part of the count rate. **B.** Concentration dependency of the aggregate formation of GFP-IDR. Increasing concentrations of GFP-IDR indicated on the top were incubated in a crowding solution in the presence or absence of 5% PEG (“PEG(+)” and “PEG(-)”). The formed aggregates were observed using a fluorescence microscope. The yellow dashed square regions are enlarged at the bottom of the figure. The scale bars in the top and bottom panels indicate 10 and 2 μm, respectively.

**Supplementary Fig. S10.**
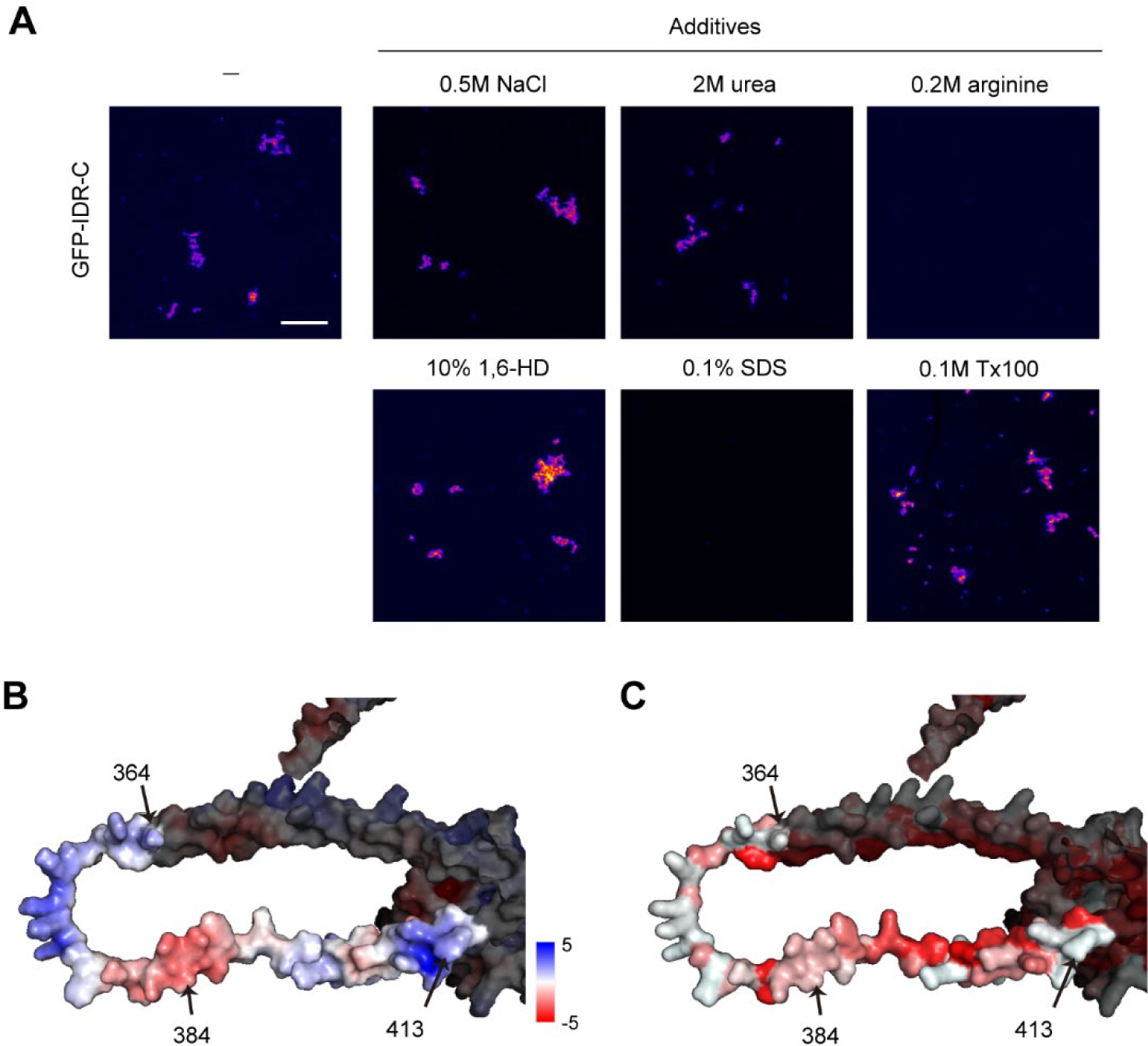
Effect of additives on GFP-IDR-C aggregate. **A.** Eight micromolar of GFP-IDR-C was incubated in a crowding solution supplied with additives indicated on the top, and the formed aggregates were observed using fluorescence microscopy. The scale bar represents 10 μm. **B.** Electrostatics of IDR. The electrostatic condition of Bqt4 was calculated by the adaptive Poisson-Boltzmann solver plugin in PyMOL software and drawn on the predicted Bqt4 structure by Alphafold2 (O60158). The blue and red portions indicate amino acid residues with positive and negative charges, respectively. Bright residues correspond to 364-413 aa. **C.** Hydrophobicity of IDR. Hydrophobicity around IDR is shown on the predicted Bqt4 structure as in (B). The red and white portions indicate hydrophobic and hydrophilic regions, respectively.

**Supplementary Fig. S11.**
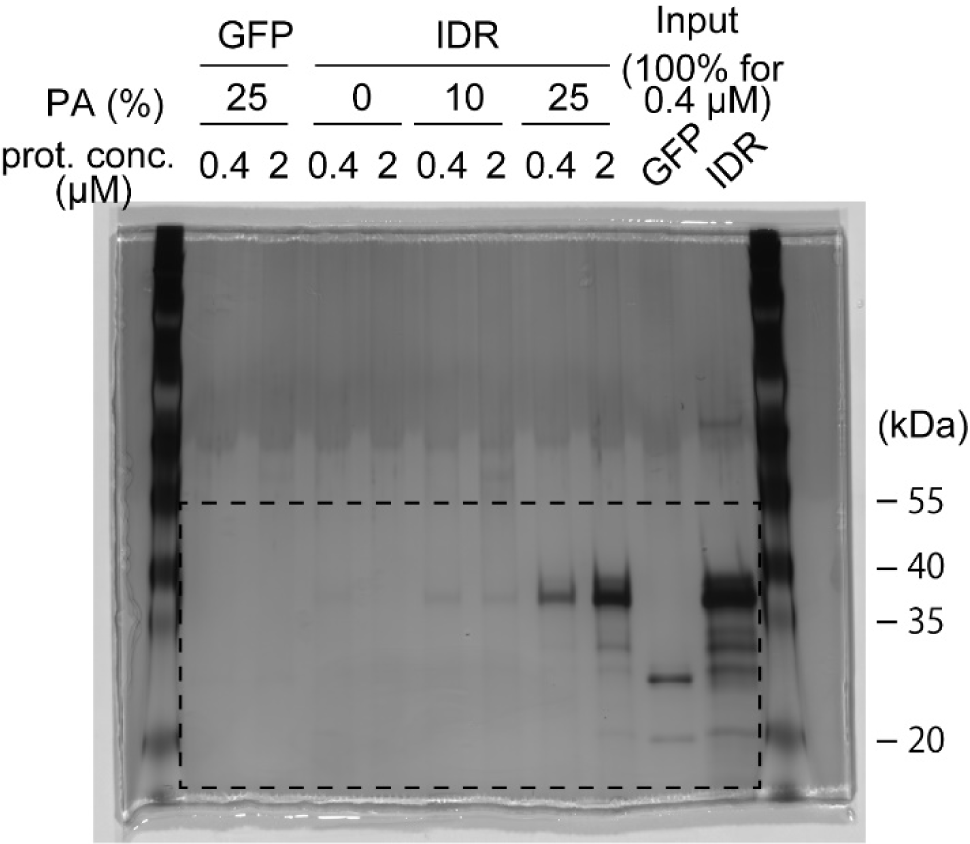
Uncropped gel image. The dashed square region is shown in Figure 4B.

## Legends to Supplementary Movie

**Supplementary Movie 1. Detachment and declustering of centromeres from the NE by Bqt4 overexpression**

GFP-tagged IDR-TM (green) was expressed under the control of the *nmt1* promoter in cells expressing Cnp1-mCherry (magenta) as a centromere marker. The cells were pre-cultured in EMMG+1 μM YAM2 for overnight, washed, and then observed using fluorescence microscopy. The bright-field image is shown on the right. The elapsed time is denoted as h:min:s. The scale bar indicates 10 μm.

**Supplementary Movie 2. Chromosome segregation defect by Bqt4 overexpression**

GFP-tagged IDR-TM (green) was expressed under control of the *nmt1* promoter in cells expressing Atb2-mCherry (magenta). The cells were pre-cultured in EMMG+1 μM YAM2 overnight, washed, and then observed using fluorescence microscopy. The elapsed time is denoted as h:min:s. Scale bar: 5 μm.

## Notes

### Competing Interest Statement

The authors have declared no competing interest.

### Summary of Updates

We corrected errors in Figure legends.

